# Temperature Sensitive Contact Modes Allosterically Gate TRPV3

**DOI:** 10.1101/2023.01.02.522497

**Authors:** Daniel Burns, Vincenzo Venditti, Davit A Potoyan

## Abstract

TRPV Ion channels are sophisticated molecular sensors designed to respond to distinct temperature thresholds. The recent surge in cryo-EM structures has provided numerous insights into the structural rearrangements accompanying their opening and closing; however, the molecular mechanisms by which TRPV channels establish precise and robust temperature sensing remain elusive. In this work we employ molecular simulations, multi-ensemble contact analysis, graph theory, and machine learning techniques to reveal the temperature-sensitive residue-residue interactions driving allostery in TRPV3. We find that groups of residues exhibiting similar temperature-dependent contact frequency profiles cluster at specific regions of the channel. The dominant mode clusters on the ankyrin repeat domain and displays a linear melting trend while others display non-linear trends. These modes describe the residue-level temperature response patterns that underlie the channel’s functional dynamics. With network analysis, we find that the community structure of the channel changes with temperature. And that a network of high centrality contacts connects distant regions of the protomer to the gate, serving as a means for the temperature-sensitive contact modes to allosterically regulate channel gating. Using a random forest model, we show that the contact states of specific temperature-sensitive modes are indeed predictive of the channel gate’s state. Supporting the physical validity of these modes and networks are several residues identified with our analyses that are reported in literature to be functionally critical. Our results offer high resolution insight into thermo-TRP channel function and demonstrate the utility of temperature-sensitive contact analysis.

## Introduction

Transient Receptor Potential Vanilloid 3 (TRPV3) is a tetrameric divalent cation channel and member of the ion channel family responsible for the molecular basis of temperature sensation[1,2]. TRPV3 is expressed in skin keratinocytes [3] and is associated with Olmsted syndrome, atopic dermatitis, and hair growth defects [4–7]. Structurally, it is characterized by a large intracellular skirt composed of ankyrin repeat domains (ARDs) (Fig. 1). Each subunit interfaces with the adjacent protomer’s ARD via an inter-protomer coupling domain (IPCD) composed of antiparallel beta strands. A transmembrane voltage sensing like domain (VSLD) and pore domain interface with adjacent subunits within the membrane. At the junction between the transmembrane and ARD domains, the namesake TRP helix lies horizontally along the inner membrane, above the so-called helix-turn-helix (HTH). Both the N and C terminal tails are disordered and almost entirely absent from available structures, however in the structures where they are resolved they exhibit different conformations depending on the channel’s state[8–10]

**Figure 1.**
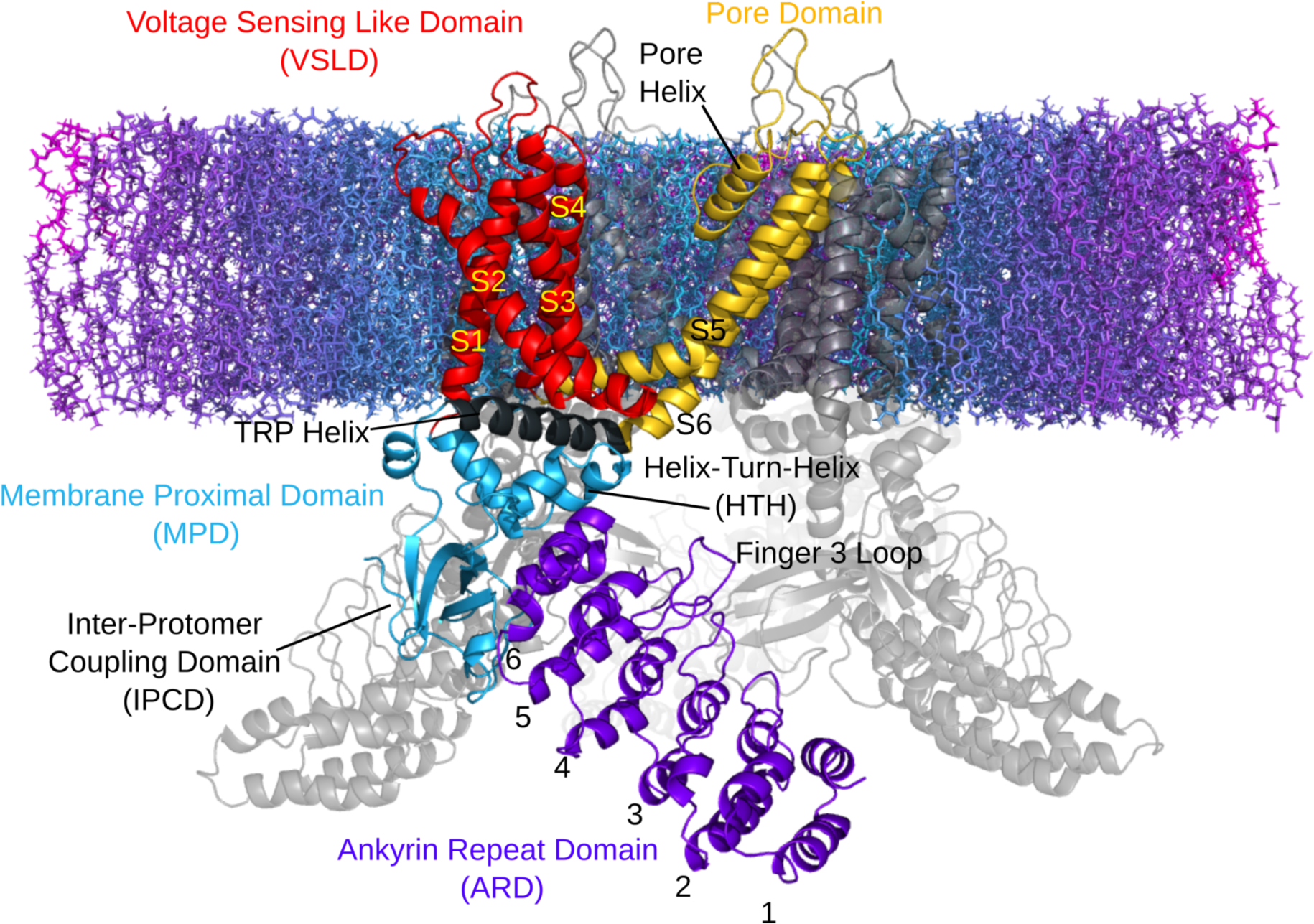
TRPV3 protomer domain structure embedded in a lipid membrane cross-section with the intracellular region below and extracellular region above.

The pore itself is characterized by a central cavity between an extracellular selectivity filter and lower gate. Upon activation, the gate radius widens by approximately 1.5 angstroms [8] to allow passage of Ca2+ ions to the intracellular space. In order to function, TRPV3 has an additional sensitization requirement where the channel must first be exposed to temperatures exceeding 50°C before it can be activated by temperatures above 30°C [8,9,11–14]. The mechanism translating temperature increase to gate expansion in thermo-TRP channels is not well understood despite a number of reported structural changes that coincide with the open gate [8,9,15–18].The identification of temperature-sensing regions and an explanation of how these regions coordinate their dynamics to control gating within a precise temperature range are of fundamental importance for understanding the design principles of temperature-sensing ion channels.

Recently, cryo-EM structures have provided a coarse description of the channel’s dynamics composed of closed, sensitized, and open states, and the inputs for high-resolution molecular dynamics simulations [8,19–27]. Despite the increasingly available structures, computational studies have yet to reveal the underlying principle of thermo-TRP regulation. In this work we employ atomistic simulations, contact network analysis and machine learning techniques to provide a mechanistic explanation for allosteric regulation of TRPV3 and identify specific regions and residues implicated in temperature sensing.

Starting from an initially closed structure of TRPV3 we use Hamiltonian Replica Exchange Molecular Dynamics (HREMD) to sample the conformational ensembles of the channel across a wide range of temperatures. Then, utilizing all-atom contact data, we identify the most temperature-sensitive inter-residue contacts via Principal Component Analysis (PCA)[28]. The application of PCA segregates the contacts into distinct modes describing their temperature-dependent behavior. These modes are centered on important regions of the channel and can be reconciled with experimental results as the residues defining them overlap with mutational and allosteric hot spots.

To identify allosteric pathways that couple domains and communicate sensation of heat across the structure, we employ network analysis methods, using contact frequencies as edge weights. Finally we use the instantaneous contact records and gate radius measurements in a random forest model to identify which temperature-sensitive contact modes best inform on the expansion of the channel’s gate. Our results identify many important interactions that are supported by previous experimental research and offer a model by which thermosensitive ion channels can exhibit activity precisely tuned to temperature. This model involves multiple structurally localized temperature-sensitive contact modes that trigger conformational changes and propagate dynamics to the gate through a network of contacts.

## Results

### Temperature sensitive contacts

Thermo-TRP channels allosterically respond to temperature changes with rearrangements that expand their pore and allow ions to diffuse through. Experimental structures provide the endpoints of this process but leave out the mechanistic details of intermediate steps. To investigate the continuous space between the endpoints, we sampled the equilibrium inter-residue contacts by simulating TRPV3 across 28 effective temperatures spanning the sub-activation, activation, and beyond. From these we calculated all of the inter-residue contact frequencies using software that accounts for different cutoff distances and interaction angles between specific chemical groups.

By applying PCA on the temperature-dependent contact frequency data [28] we extract principal components which describe the ion channel’s collective contact making and breaking responses to temperature. Each contact has a unique loading score (eigenvector component) describing the contact’s contribution to the variance described by the PC. Here the normalized loading score is used to rank a contact’s temperature-sensitivity within different PC trends, or “modes”. These modes can reveal where temperature-sensing function is localized as well as potential allosteric connections since the loading scores within a PC are a measure of correlation between contact responses.

We utilized the difference of roots test [29] to determine the number of PCs to include in further analysis (Fig. 2A). By taking the difference of the explained variances between adjacent eigenvalues, it’s possible to identify where the change in explained variance between the original PCs drops below the change in explained variance exhibited by the null hypothesis data (no covariance between contact frequencies across temperature). This test identified the first 7 PCs as significant for describing the structure’s temperature-dependent contact behavior. Though the difference in explained variance drops back below the threshold for PC9, the trend is clearly broken after PC7 and indicates that the subsequent PCs are becoming noise.

**Figure 2.**
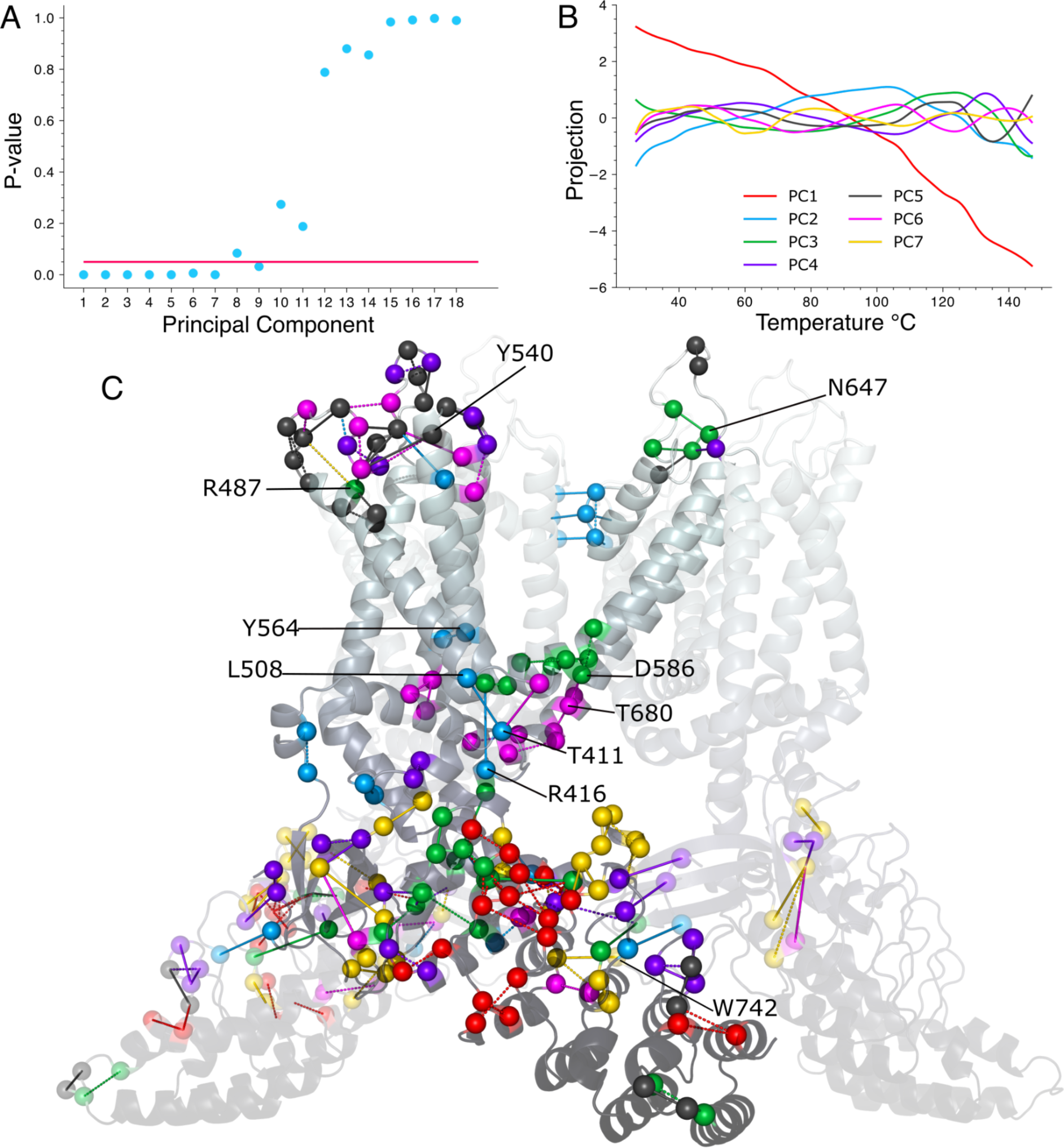
Temperature-sensitive contact analysis of TRPV3. **A.** Difference of roots significance test. Horizontal line indicates a P value of 0.05. 18 of the 28 PCs shown. **B.** PC projections showing the general trend in contact frequencies across temperature for the first 7 PCs. **C.** The most temperature-sensitive contacts (top 18) from each of the top 7 PCs. The contact is colored according to the PC on which it has the highest loading score (rank) following the same PC color scheme in B. Solid lines indicate contacts that tend to increase in frequency within the activation temperature range and dashed lines indicate contacts that tend to decrease in frequency (based on linear fits) in the activation temperature range. The contacts surrounding the IPCD are duplicated on either side of the central protomer as are those extending down to the distal end of the left protomer’s ARD. Allosterically important residues discussed in the text are indicated.

We hypothesize that these contact modes are a fundamental mechanism for TRPV3 to sense and initiate a response to temperature. Before we can connect them to the gate’s function, we first identify where the temperature-dependent contact modes are localized, which residue pair interactions best define the modes, and what temperature-dependent patterns their contact frequencies exhibit.

The PC1 mode accounts for nearly 80% of the variance in the contact frequency data (Fig S1) and captures the expected melting trend where contact frequency decreases with increasing temperature (Fig 2B red). PC2 accounts for roughly 8% of the explained variance and describes a less intuitive consequence of increased temperature: an increase in contact frequency for many of its highest ranking contacts (Fig. 2B blue, see S2 for examples of individual contact frequency plots). The subsequent PCs describe trends that oscillate within progressively narrower temperature and frequency ranges. By PCs 5-7 a local maximum amplitude develops around the sensitization temperature of TRPV3 (fig 2B vertical dashed bar).

### Structurally Clustered Modes Identify Temperature Sensing Regions

Structural clustering is evident among the top ranking contacts of the first seven PCs (Figure 2C). Many of the most temperature-sensitive contacts in these clusters have experimental significance that we note here. From thousands of contacts, we selected the 18 highest ranked contacts on each PC to represent the highly temperature-sensitive interactions as these all fall on the upper half of the narrow range of rapidly decaying loading scores (Fig S3, see SI Table 1 for loading score rankings). Thus the following correspondences between these contacts and published data on functionally important residues is remarkable and evidence that these modes have allosteric significance.

Regarding the thermodynamics of TRP gating, Yao et al. showed that ankyrin repeats 5 and 6 were responsible for the activation enthalpy differences between TRPV1 and TRPV2 [30]. This observation can easily be explained in the context of TRPV3 by the densely clustered PC1 melting trend that spreads out from ankyrin repeat 5 (Fig 2C red). Indeed, PC1’s prominent role on the ARD becomes more apparent when expanding the range of the depicted contacts’ ranks(Fig. S4).

In contrast to PC1, most of PC2’s top contacts display increasing frequency with temperature (across the natural activation temperature range). An expanded set of PC2 contacts clusters heavily around the HTH and MPD, overlapping with PC1 on the upper ARD and extending into the VSLD and inter-subunit contacts on the IPCD (Fig S5). In the context of the first 7 PCs’, PC2 best describes a dispersed set of generally order promoting interactions mostly surrounding the TRP helix (fig 2 C blue, solid lines). Perhaps the most notable correspondence with experimental work is the overall highest ranked PC2 contact between T411 on the HTH and L508 on the S2-S3 linker. T411 is at (or in terms of substitutions, “adjacent to”) the position identified by Liu and Qin[11] where an insertion of a cognate residue from TRPV1 eliminates the use-dependent sensitization of TRPV3. In fact, both residues of this contact carry significance as a recent study showed L508 to be the most important residue in binding the allosteric activator carvacrol [31]. PC2’s 10th ranked contact involving Y564 and F522, which connects the S4 and S3 helices, also has experimental significance. When mutated to alanine, Y564A enhances binding of the allosteric activator 2-APB [18,32], a ligand which has played an important role in TRPV3 studies.

PC3 (Fig. 2C green) describes a mode with contacts localized mostly to repeat 6 on the upper ARD, adjacent to the PC1 cluster, and the lower transmembrane region at the S5 bend. Though structurally isolated, PC3’s 3rd ranked and most notable experimentally relevant contact is on the S5-Pore helix linker between N647 and C619, as an N647Y mutation was shown to eliminate activity[33].

PC4 clusters on both the extracellular loops and around the IPCD (Fig. 2C purple). Because these regions are so distant, it’s likely that PC4 encompasses the dynamics of structurally independent modes that exhibit similar temperature-dependent contact frequency profiles. PC5 is the dominant mode of the extracellular VSLD loops (Fig. 2C dark grey) where most of the contacts display a subtle increase in frequency within the activation temperature range. Residue Y540 is involved in the 7th and 9th most temperature-sensitive contacts on PC5 and 6 respectively and is involved in a binding site of the allosteric activator 2-APB [32] as is the nearby residue R487 that participates in multiple modes (rank 4 on PC3, rank 9 on PC5).

PC6 clusters heavily along the TRP helix/S6 bend (Fig. 2C magenta) just below the gate itself, precisely where structural changes are associated with gating [32]. Most notable on PC6 is the 5th ranked, functionally critical contact between T680 and D586. Mutating T680 to alanine to break its hydrogen bond with D586 abrogated activity in an experimental TRPV3 construct [15]. These PC6 contacts also surround residues 568, 573, and 692 at the S4-S5 bend and TRP region where mutations are implicated in Olmsted Syndrome [4,5].

Lastly, PC 7 clusters heavily around the IPCD and adjacent finger 3 loop(Fig 2C gold). Several of the residues in PC7’s highly temperature-sensitive contacts surrounding the IPCD are involved in the state-dependent exchange of the N and C terminal tails[8] including R226, N273, K743, and H745. Among the highest ranking PC7 contacts (rank 6), W742 was shown to substantially reduce channel activity upon mutation to alanine [9].

Overall the correspondence with experimental data presented here confirms that these contact modes are capturing a functionally relevant phenomenon. Interestingly, the lower eigenvalue PC projections (4-7) show a local maximum (Fig 2B vertical dashed line) in the temperature range where activation of TRPV3 occurs [11]. Since PC6 is clustered immediately by the gate, we found it likely that the overlapping activation maximum was not merely a coincidence. However, the other PC clusters (which exhibit local maximums in their trends around activation) are quite distant from the gate, prompting us to investigate a means of communication between them and the gate. We hypothesize that these contact modes influence the channel’s motions and that the motions regulate gate expansion by transmitting dynamics through an allosteric network.

### The Motions of the TRPV3 Protomer

The contact frequency PCs do not directly inform on the 3-dimensional motions that the channel samples but the making and breaking of contacts certainly influences them. Since cryo-em structures confirm that distributed conformational changes correlate to the state of the gate, we sought to relate the channel protomer’s contact modes to its motions. To this end we performed PCA on the protomers’ Cartesian coordinates from the 54 °C simulation (simulation temperature immediately above the sensitization threshold) and extracted the endpoints of the largest motions from the first 6 PCs (Fig. 3) which describe from 5 to 20% of the variance (50% total) (Fig. S6).

**Figure 3.**
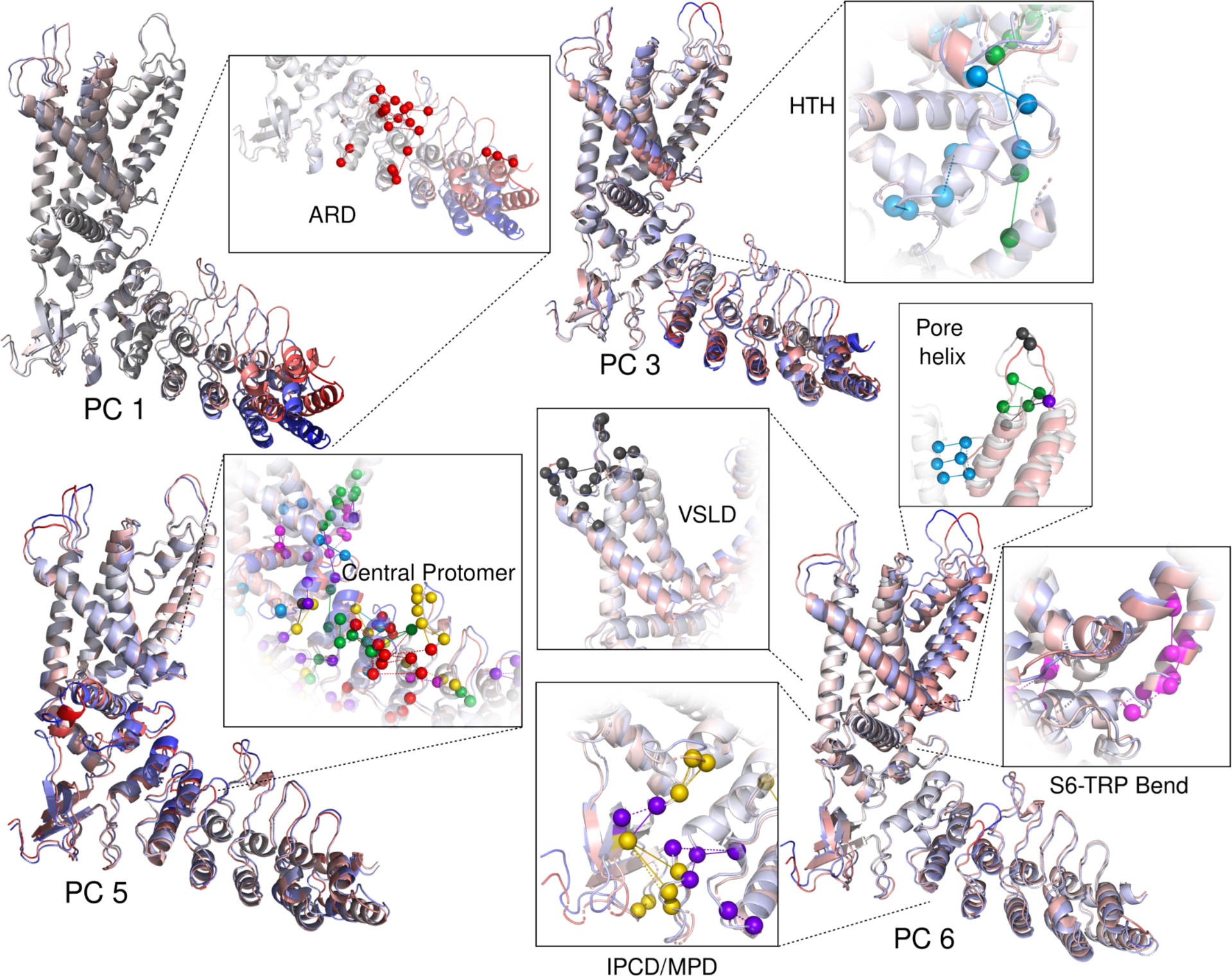
Cartesian principal components and temperature-sensitive contacts. Extremes of the protomer structures (red or blue) extracted from the corresponding cartesian PCA trajectory projections and colored according to regions of maximum displacement. Insets show highly temperature-sensitive contacts implicated in the motions of specific regions.

The fact that the large bending and twisting motions of the ARD seen on Cartesian PCs 1 and 2 (S7) originate next to the contact PC1 cluster suggests that these dynamics and contacts are related (Fig 3 PC1 Inset). The melting of contact PC1 apparently creates a hinge in the middle of the ARD, allowing the lower ARD to swing.

Cartesian PCs 3 and 5 show the HTH, TRP helix and S3-S4 bend compressing together at one end of the concerted motion implicating the order promoting interactions of PC 2 and 3’s contacts around the HTH (Fig 3 PCs 3 and 5 insets, blue and green spheres). Notable also is the correlation between the extracellular loop motions (particularly those on the VSLD linkers) and the motions of the transmembrane helices. The extracellular loops here are dominated by contact PC 5, making their interactions the likeliest source of these upper transmembrane dynamics (fig 3 PC6 upper left inset, black spheres). The subtle IPCD motions on several Cartesian PCs are most attributable to the interactions described by contact PCs 4 and 7 as they surround the IPCD (Fig. 3 lower left inset). Naturally, the motions of the S6-TRP bend are most attributable to contact PC6 (Fig. 3 lower right inset).

The motions of the first 5 cartesian PCs occur in regions of the channel distant from the gate while the 6th PC captures the largest dynamic of the pore domain. Of the motions shown in Figure 3, the 6th cartesian PC is the most likely to capture gate expansion. We noticed that on this cartesian PC, the S5 and S6 motion is correlated with motions at the distal tip of the ARD and the IPCD. Whether or not this suggests that one is influencing the other, if the temperature-dependent contact dynamics regulate gating, then there must be a means of transmitting force (or more generally, “dynamics”) generated by the motions of various regions to the gate. Network analysis methods lend themselves here as a means of identifying signal transmission pathways.

### Allosteric networks couple domain motions in TRPV3

Though debate regarding the presence of a single vs. distributed temperature-sensing module has not been fully resolved, experiments on several TRP channels suggest that a single domain is responsible for determining the gross activation temperature range [34],[30,35] with the coordination of multiple domains necessary for gate activation and fine-tuned temperature specificity[33,36,37]. A prevalent view is that these multiple regions respond by enhancing inter-domain ‘coupling’[9,38–41]. Here we perform network analysis to investigate the temperature-dependent domain coupling by applying the contact frequencies as network edge weights. We use data from the 27°C simulation to represent the unactivated channel’s network as it is our lowest temperature simulation (and below it’s physically active temperature) and the 54°C simulation data to represent the activated channel as it is the first temperature in our simulation ensemble that exceeds the >50°C initial sensitization requirement [18] and is within the temperature range of near maximum experimental recorded activity[11].

Network analysis techniques have been used to identify communication pathways in proteins as well as communities of residues that comprise functional domains [42–46] and can thus offer a description of domain coupling. Indeed, similar analysis has been applied to coarse-grain simulations of TRPV1 [19].

### The Network Community Structure of a TRPV3 Protomer

A network can be partitioned into communities with the Girvan-Newman [47]algorithm by calculating all of the shortest paths (based on contact frequency edge weights) between nodes (residues, in the context of proteins), removing the highest betweenness centrality[48] edge and then repeating this process until the network has been broken up into a pre-defined number of communities with no more edges connecting them to one another.

We initially partitioned the TRPV3 protomer into 5 communities which returned the anticipated community structure that corresponded very closely to the accepted domain structure; however by specifying 6 communities we discovered that the community structure displayed a visible correspondence with the dynamics and temperature-sensitive contacts of the ARD. Specifically, a clean division of the ARD community structure occurs (Fig. 4A gold and purple communities) at the region in which both our simulations (Fig. 3 PC1, S7 PCs 1 & 2) and earlier published coarse-grained simulations on TRPV1[19], as well as structural data [8,18,32] show large motions develop on the first repeats. This boundary of the two ARD communities is also conspicuously located directly under the disordered tail binding site and IPCD (see PDB 7MIO) [49], adjacent to the finger 3 loop [50],and on the edge of the contact PC1 cluster. Thus we proceeded to investigate the community temperature dependence in terms of 6 communities.

**Figure 4.**
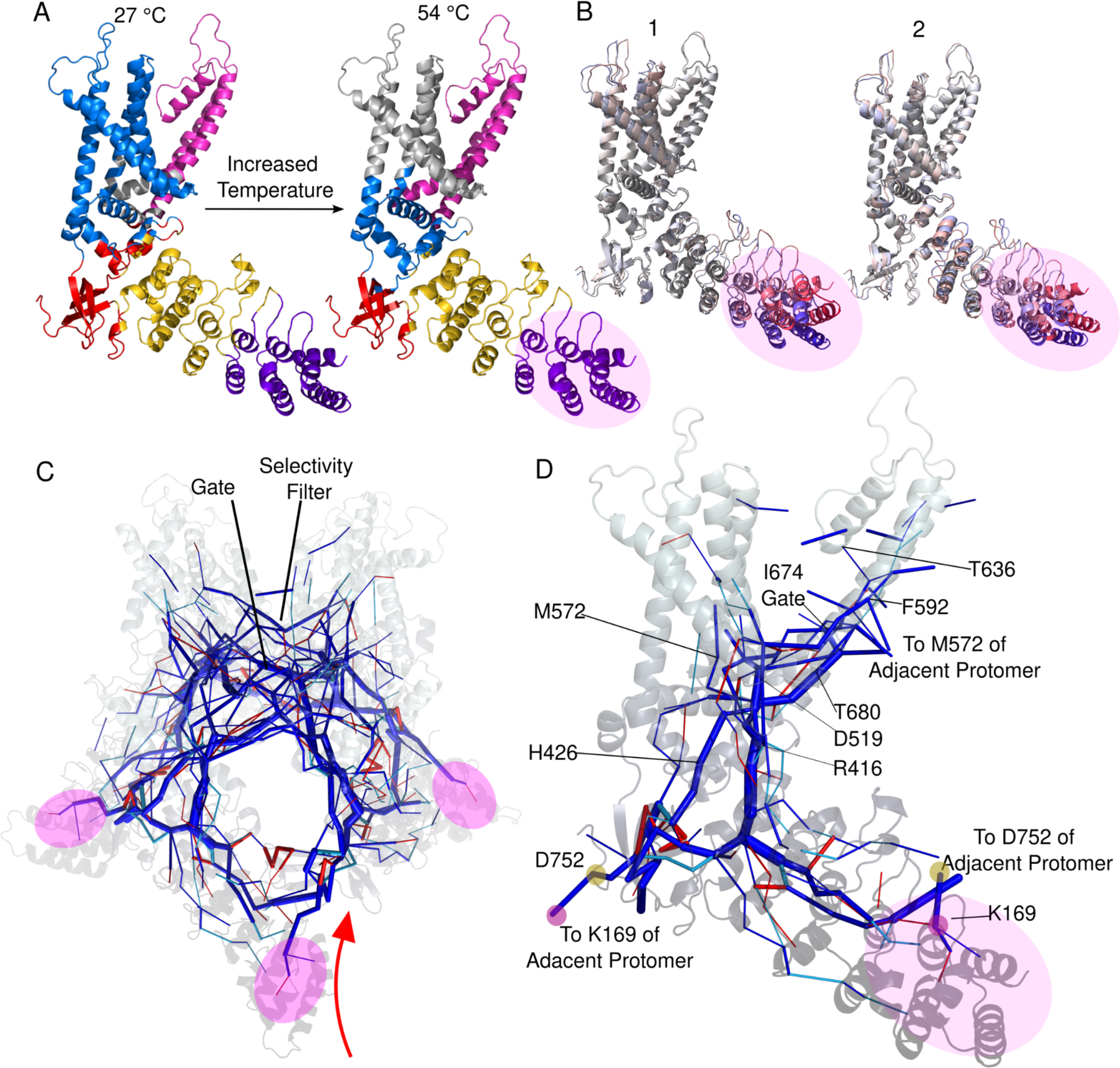
Network analysis of a TRPV3 protomer based on average contact frequency edge weights. **A.** Community structure of a TRPV3 protomer changes from low to high temperature **B.** Cartesian PC 1 and 2 projections showing the dominant motions occur on the distal ARD. The dynamic region corresponds to the purple community in A and is highlighted on the network in C and D. **C.** Tetramer’s “allosteric skeleton”. Thickness of the line indicates magnitude of betweenness centrality. Dark blue edges exist at both 27°C and 54°C. Red edges are present in the top centrality edges at 27°C but not at 54°C. Light blue edges occur in the top edges at 54°C but not 27°C. **D.** Individual protomer’s allosteric skeleton with residues discussed in the text that lie on the network pathways.

### Temperature Dependent Coupling of Network Communities

Since temperature is proposed to enhance coupling between domains, leading to channel activation [38], we expected the community structure to change with temperature. While the ARD maintains almost identical community structure between temperatures (Fig. 3A purple and gold communities), this is not the case for other critical regions of the protein. At high temperature the S6 community (magenta), which contains the gate, extends into the S6-TRP bend and pre-S5 region around the central protomer. The HTH also exhibits a change in its community structure, albeit a subtle one considering that the HTH is composed of few residues. The residues composing the HTH are distributed among the IPCD, ARD, and VSLD communities at low temperature but participate largely in the same community defining the TRP helix and pre-S1 at high temperature. The HTH serves as the intersection of other communities including those of the S3-S4 kink (via T411 at both low (blue) and high temperatures (grey)), TRP helix (blue), and ARD (gold), reflecting its central position in the structure’s network.

Thus, community structure changes indicate that there is an enhanced coupling between the intracellular HTH and the lower transmembrane region of the VSLD and between the pore domain and region adjacent to the central protomer. Given this evidence of altered domain coupling around the central protomer, we sought to identify the precise pathways by which the intracellular and extracellular dynamics can be transmitted and integrated to regulate the gate.

### Edge Betweenness Centrality Identifies Allosteric Pathways

The TRPV3 selectivity filter is constitutively active and apparently not under allosteric regulation of other domains [39]. Because the gate around residue I674 is the only other constriction point within the pore, allosterically important dynamics must somehow be integrated here to precisely regulate a ∼1.5 angstrom expansion. To investigate the pathways by which the distant domains can communicate with the gate, we calculated edge betweenness centralities using contact frequencies from the 54°C simulation (where a network associated with the sensitized /active state should exist) as edge weights. We also calculated the network from the 27°C simulation to identify where temperature-dependent changes in the network might occur. Edge betweenness centrality measures how many shortest paths between all other residues in the network pass through a given edge (contact). Figure 3B shows the most central edges (Fig. S8, left of vertical red cutoff bar) on the protomer structure with clear pathways running from the distal region of the ARD through the HTH and branching into the VSLD and pore domain. Multiple pathways from distant regions converge on the gate residue of I674 and the adjacent S5 helix whose motions can sterically influence the gate.

Several observations support that this “allosteric skeleton” includes physically real communication pathways as ligand binding sites and function-affecting mutation positions are among the residues defining the highest betweenness centrality edges. For instance, T636’s contact with Y594 is the primary edge between the Pore and S5 helices and is directly adjacent to the S6 pi bulge [51]. Mutating T636 to serine resulted in a hyperactive mutant [52] and this same position is known to be a binding site for the inhibitor scutellarin [53].

A highly central inter-protomer edge passes through M572 and F592, between the PC6 cluster and region associated with Olmsted Syndrome. Given that a major inter-protomer communication pathway and temperature response (PC6) exists here, it might be expected that mutations to this region would have functional consequences.

The interaction between H426 and R693 is a central edge on the most direct path connecting the IPCD and the TRP helix. H426 binds the agonist 2-APB [54] and both residues participate in binding the allosteric inhibitor Osthole [49]. The functionally critical contact between T680 and D586 (mentioned above) is both a top temperature-sensitive contact and highly central edge suggesting it is of particular allosteric importance [15]. The contact between R416 and D519 connects the HTH to VSLD and is one of the only other contacts that is both highly temperature-sensitive and a highly central edge.

Qin and colleagues [11,55] have shown the large impact of mutations on this region and given that the HTH is the most direct route connecting a protomer’s ARD to the transmembrane domain, this contact and and the aforementioned T411-L508 are likely critical for domain coupling as they both increase in frequency with temperature.

Interestingly, the most central edges from the distal ARD (repeats 1 and 2) pass through the IPCD to the adjacent protomer, rather than down the originating protomer’s ARD. This suggests that dynamics at the distal ARD might be most effectively transferred through the IPCD to influence the gate. Additionally, the most central edge connecting the two protomers here is between K169 and D752, a salt bridge known to be critical for function[9]. With only the edges depicted in figure 4D, an unbroken path can be traced from K169 on the adjacent protomer through the contact PC6 cluster on the TRP bend and to the gate residue of I674. These pathways indicate that there is indeed a means for the dynamics caused by making and breaking of temperature-sensitive contacts to allosterically communicate with the gate. We were thus compelled to search for a quantitative link between the temperature-sensitive contact modes and the dynamics of the central gate.

### Evidence of Allosteric Gating Regulated by Specific Contact Modes

To investigate the connection between the contact frequency modes and the regulation of the gate, we turned to supervised shallow machine learning, specifically a random forest, as it can pull important features out of the inherently noisy biomolecular interaction space and provide readily interpretable model coefficients.

Here we utilized the Hole [56] program to determine the gate radius (Fig 5 A & B) and construct binary labels to classify the gate at each frame in the trajectory. Hole uses an algorithm to identify the best route to pass a sphere of variable radius through the channel pore. As the sphere’s radius is altered to avoid overlap with atoms lining the pore, the pore radius is measured. These calculations revealed that the gate did not sample a fully open state during the course of the simulation, perhaps due to the missing tails which are known to be important for activity[8,9,37]. In the presence of an approximately -70mV membrane potential, Ions can pass through channels at a rate of up to 108 per second [57,58]. Based on the work on TRPV1[59] a lower estimate of 2 107 per second is reasonable at 50°C. If the channel were to open at activation temperature in the absence of a membrane potential at the same rate as ion conductance, we would have expected at least 2 open observations during our simulation.

**Figure 5.**
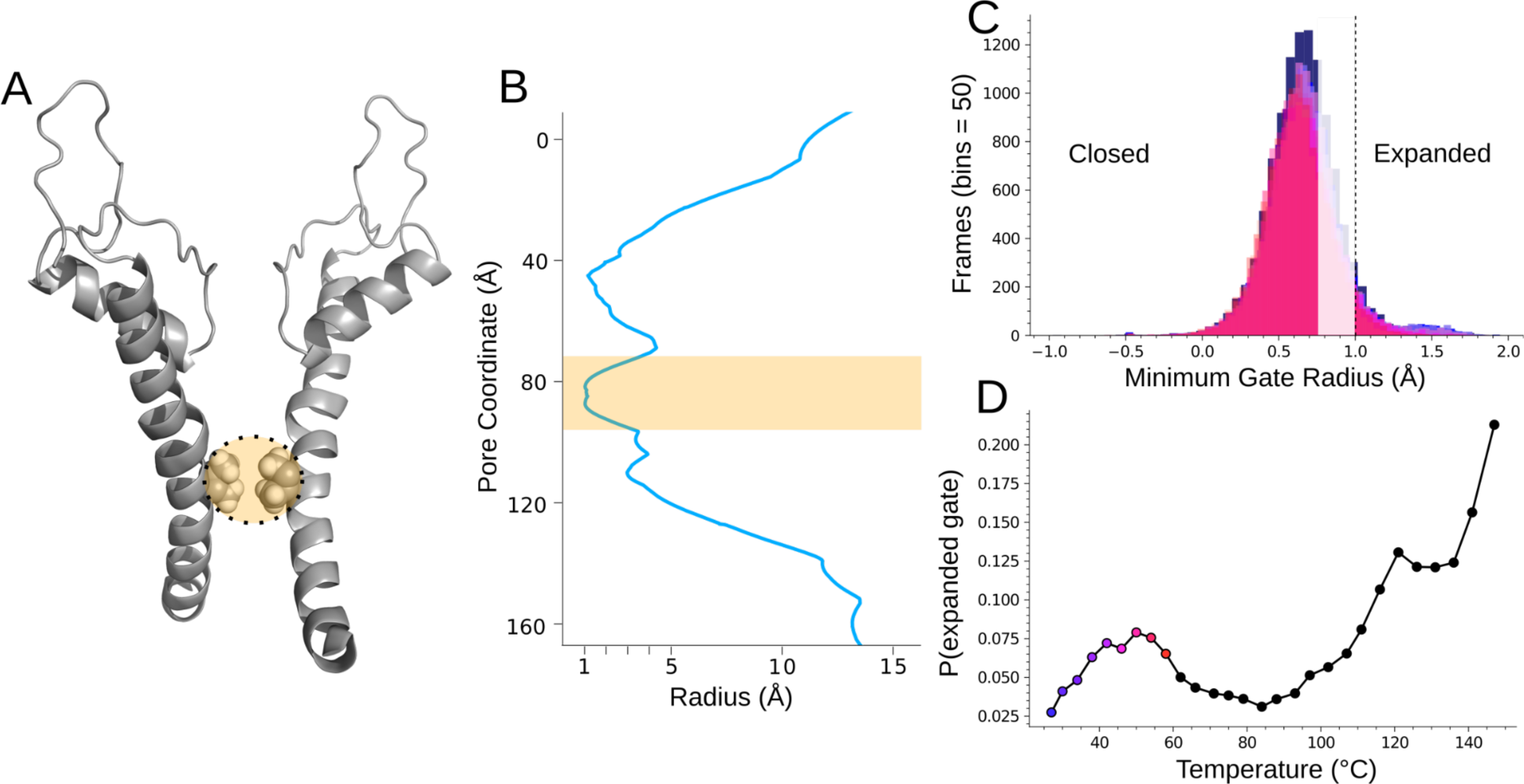
Gate measurement. **A.** Two opposing S6 helices defining the pore region of TRPV3 with I674 gate residues highlighted. **B.** Example Hole analysis pore profile with the gate region highlighted. The minimum value in the highlighted region was used to define the state of the gate for the random forest model. **C.** Overlapping histograms of the gate radius from the 8 simulations used to construct the random forest input data. The faded region of the histogram represents the gate data that was not included in the random forest in order to discretize the expanded and closed states. **D.** Probability of the expanded gate vs. temperature showing a local maximum around sensitization temperature. Colored dots indicate the simulations from which the random forest data was extracted.

Since the channel likely remains open at elevated temperature to allow diffusion down the ion’s concentration gradient, it is even more likely that, if the channel construct was behaving as a sensitized full-length construct, a fully open state would have been observed. Beyond the issue of the missing terminal tails, we speculate that the lipid membrane remaining at 27°C, due to the Hamiltonian treatment being exclusively applied to the protein, may have also prevented wider channel expansion. As such, we defined an “expanded” gate as a minimum radius greater than 1 angstrom (Fig 5C) (corresponding to the radius of a desolvated Ca2+ ion [60]) and calculated the number of frames at each temperature where the radius was above the cutoff. While the channel would need to expand by another 1-1.5 angstroms to convey a solvated Ca^2+^, our cutoff represents conformations significantly more open than the equilibrium closed radius captured by cryo-em. Figure 5D shows that the P(expanded gate) (number of frames where gate radius > 1 Å/ total frames) trends up from baseline near the physiological activation temperature to a local maximum at sensitization temperature (before curiously returning to baseline and only spiking again after ∼100°C) indicating that a dynamic relevant to channel opening is occurring in the temperature range where the channel is naturally activated.

The sample data for our random forest model was the binary state of each contact at each frame of the simulations. With this, we trained a random forest to identify an expanded gate based on the contact states of a TRPV3 protomer. To briefly summarize, a random forest is composed of “decision trees”. Each tree is trained using a random subset of contacts. The data is split at each node (branch) according to the features (contacts) that best separate the data into the two classes (closed or expanded gate). For inference, the random forest uses the prediction made by most of its trees. The individual contacts that provide the most information on the gate’s state and thus split the data most effectively, are assigned high feature importance values during the fitting process. The purpose of building this model was to extract the contacts to which the model assigned high feature importance, revealing the interactions and corresponding modes that influence gate expansion.

We fit several models with various parameters that all returned approximately 65% recall and nearly 90% precision at predicting the expanded channel state. From two of these models we took the contacts with feature importances above .001. Figure 6A shows that beyond these first roughly 100 contacts, the feature importance drops to essentially 0 and the remaining contacts become uninformative. The feature importance is normalized and thus dependent on the number of features included in the model. This cutoff value was determined by plotting the feature importance vs the corresponding contact in descending order which reveals that only a small subset of contacts are relevant to predicting the state of the gate (Fig 6A). We then determined on which PCs these feature important contacts have their highest loading score. We took the intersection of the most feature important contacts (>.001) which returned 106 shared contacts and found that PCs 1,3, are underrepresented among the predictive contacts while PC 2 and the lower eigenvalue PC contacts occur more frequently (Fig 6B). Considering that PC1 accounts for the overwhelming majority of the variance in contact frequency, it is noteworthy that the predictive contacts rank highest on other PCs and further indicates that these contact modes capture a functionally relevant phenomenon.

**Figure 6.**
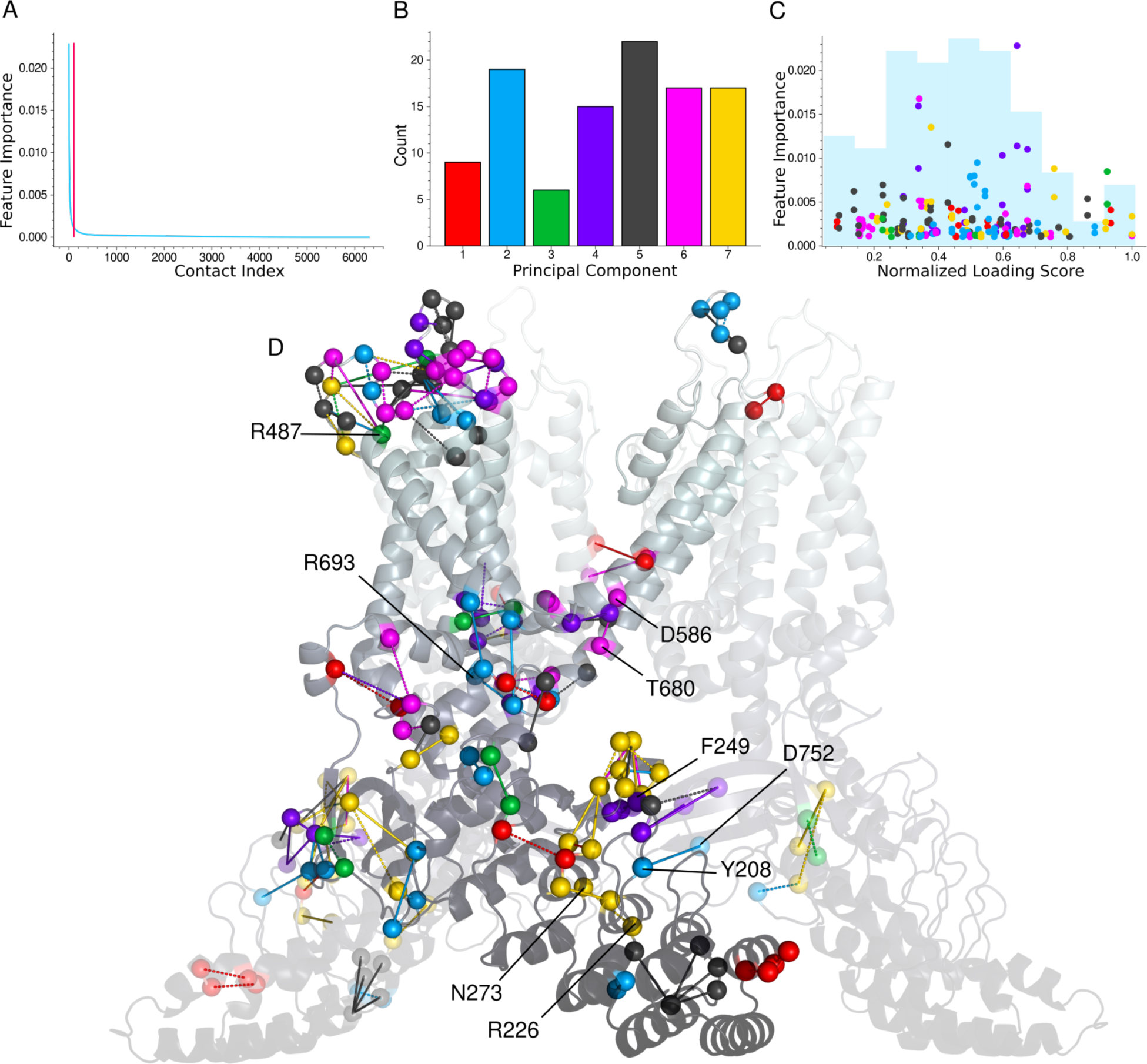
Random forest feature importance analysis. A. Sorted feature importance scores showing a small minority of contacts contribute to predicting the gate’s state. Only contacts left of the vertical cutoff bar are included in the remaining analysis. B. The counts of each PC’s contacts that occurred in the intersection of the 2 random forests’ top shared feature important contacts. C. Feature importance vs. temperature-sensitivity for the same contacts. The contacts were assigned the PC on which they have their highest loading score and colored accordingly. The pairs of vertically aligned dots appear because feature importance values from both random forests are shown. Histogram of loading scores in faint blue. D. Depiction of the feature important contacts on the TRPV3 protomer with the contacts on the ARD and surrounding the IPCD duplicated to illustrate their influence from the adjacent subunits.

The structure depicting the high feature importance contacts (Fig 6D) is reminiscent of the conformational wave which accompanies state transitions proposed by Nadezhdin and coworkers[8]. The histogram in the background of Figure 6C indicates that the contacts most predictive of the expanded gate occur not only primarily amongst the lower eigenvalue modes (6B) but also in the middle of the temperature sensitivity scale. Although structural clustering of contact modes is less apparent than when considering only the top temperature-sensitivity ranks, the distribution of loading scores and PCs among the predictive contacts illustrates the channel function’s reliance on many interdependent dynamics. This suggests that allosteric trigger points might be localized (temperature-sensitive contact clusters), but the transmission of allosteric signals might be distributed more broadly, perhaps taking several of the high betweenness centrality routes. Among these predictive contacts, residues implicated in the state-dependent tail binding around the IPCD occur including Y208, R226, F249, N273, and D752[8]. F249 was shown in a 2013 study to have a significant impact on activity upon mutation to glycine[50]. Also, the contact between D586 and T680 appears along with the Osthole binding residues R693 and R487 are found. R487 has the highest feature importance (12th overall) of the experimentally significant residues. Interestingly 2 of the 3 most feature important contacts involve the truncated construct’s C-terminal residue I756.

The functional significance of the PC7 IPCD clustering in both feature importance and temperature-sensitivity is supported not only by the state-dependent exchange of the disordered terminal tails around the IPCD [8,9] but also by a 2003 study showing that the distal end of the TRPV1 C-terminal tail confers thermal specificity [37]. The small magnitude of P(expanded gate) appears to recapitulate the reduction in heat-evoked currents seen with their 72 aa truncation, given that the simulated structure lacks a similar number of C-terminal residues. In line with these observations, indicating a correlation between the IPCD and gating, is that PC7’s highest ranking temperature-sensitive contacts show contact frequency changes of similar magnitude to P(expanded gate) in the activation temperature range (Fig S9).

PC7 also exhibits the most highly ranked temperature-sensitive contacts among the feature important contacts (6C gold dots cluster at right end of plot). This is the least responsive mode of the 7 contact PCs investigated and the fact that the feature important contacts from higher eigenvalue PCs tend to have moderate to low temperature-sensitivity rankings and that PC1 is under-represented among the feature importance implies that the gating of the channel is not directly triggered by the most responsive contact modes, rather that it is buffered.

## DISCUSSION

Despite decades of work on temperature sensing ion channels, a detailed description of how they sense distinct temperature thresholds and allosterically regulate their pores in response remains lacking. In this work we employed molecular simulations, temperature-sensitive contact analysis, graph theory, and machine learning to reveal details of allosteric regulation in TRPV3.

We used temperature-sensitive contact analysis to decompose the channel’s temperature response into several structurally localized, collective residue interaction modes and show that they likely underlie the channel’s dynamics. Our network analyses describe a temperature-dependent domain coupling and reveal a network of high centrality contacts that connects distant regions of the protomer to points on and directly adjacent to the gate and appears to convey dynamics of the distal ARD through the adjacent protomer. The random forest model reveals correlations between specific temperature-sensitive contact modes and gate expansion, and points to regions of the channel whose dynamics may regulate the final step in gating.

The results offer a model for TRPV3 function that accounts for published observations and suggests a means of specific temperature tuning for other TRP channels. First, the overwhelming majority of the melting described by contact PC1 develops adjacent to the dynamic lower ARD, suggesting that it influences these motions. The ARD’s motions transmit force and alter contact states through the allosteric network, likely contributing to gate activation. Indeed, a precedent for this mechanism exists in another TRP member, the NOMPC mechanotransduction channel, whose large ankyrin repeats convey force that underlies its gating[61]. The PC1 mode also partly explains the channel’s change in heat capacity related to activation which requires the exposure of roughly 20 nonpolar residues to solvent [62,63]. The entropic payment for the increased local order described by the lower eigenvalue PCs can also be attributed to it. Moreover, the ARD is known to be the primary temperature sensor for (rattlesnake) TRPA1 [35] and, given the high explained variance of PC1, it appears to serve the same role here. Thus the PC1 mode promotes disorder which evolves with temperature to a point where numerous distributed interactions form, following the progressively more subtle PC trends, before the probability of channel opening is high enough to be physiologically relevant.

While the ARD dynamics influence the channel from the intracellular side, the VSLD loops influence it from the extracellular side. The structural clustering of contact PC5 in particular makes it the most apparent source of the subtle bends and shifts of the VSLD helices which must sterically influence the gate, and based on their projected trends, their interactions appear grossly tuned to the active temperature range of the channel.

The random forest suggests that the lower eigenvalue PC modes exert the most influence on gate expansion. Of the first three PCs, the random forest found only PC2 to have a number of contacts similar to the last four PCs that are predictive of the expanded gate. PC 2’s strong order promoting trend, clustered in the middle of the protomer, must then be critical for the integrating the various domains’ dynamics and reorganizing the allosteric network that allows the lower eigenvalue mode dynamics originating from the intracellular region to communicate with the gate.

The lower eigenvalue modes necessarily display the weakest temperature response of the PCs investigated but, as a consequence, offer an explanation for the robust regulation of the different thermo-TRP activation temperatures. A tightly controlled activation mechanism producing a small ∼1.5 angstrom expansion concomitant with a slight helix reorientation should naturally be encoded by the subtler trends such as PC 6 which clusters near the gate and PC7 which clusters around the IPCD, regions displaying state-dependent conformations. Once the channel is primed for activation by heat-induced contact mode evolution and network reorganization, a subtle dynamic originating around the highly predictive contacts in the vicinity of the IPCD apparently regulates the final step in gate expansion.

By burying functionally critical dynamics in lower eigenvalue modes and in the middle of the temperature-sensitivity scale, activation is effectively buffered from noise; an essential feature for biomolecules designed to report on noxious stimuli. From a design standpoint, controlling function with several “layers” of modes offers nature (or protein engineers) multiple “dials” with which to precisely tune function.

The largest magnitude dynamics occurring at the distal end of the ARD are transmitted most directly via the IPCD to the gate. These observations underline the importance of the disordered tails and their relationship with this region. It’s possible that their interactions here enhance dynamics transmitted to the pore region and perhaps stabilize the channel in an open state. Indeed the entropic force generated by disordered tails can tune activity [64] and the limited gate expansion we observed along with the predictive significance of the terminal contacts might be best explained in light of the missing terminal tails and the additional forces and possible network stabilization they could provide. Future computational studies should concentrate here.

Along with the description of allosteric regulation of TRPV3 presented here, we have introduced temperature-sensitive contact analysis as a framework for investigating other heat sensitive ion channels. Further, investigation of biomolecular dynamics using these modes need not be restricted to phenomena transecting temperature, as they describe correlated interactions that conceivably exist at any temperature where the protein maintains its native fold. As we have shown here, with the right reaction coordinate, machine learning techniques can be used to map these modes to their functional roles.

## Methods

### Simulations

Using the CHARMM GUI [65,66] Membrane Builder tool [67–69] and CHARMM36 forcefield[70], we assembled our simulation system around the closed PDB 7MIJ structure[8] with which we could capture processes that occur as it moved up through temperature. The 7mij structure is missing its disordered tails but importantly, there are no missing loops that require modeling. We formulated the membrane composition similar to the system simulated in Nadezhdin et al. [8] with the lipid models available in in Charmm-Gui as follows: 28% cholesterol, 13% phosphatidylglycerol, 22% phosphatidylcholine, 13% phosphatidylethanolamine, 4% phosphatidylinositol, 9% sphingomyelin, and 13% phosphatidylserine. Ionizable residues were protonated at neutral pH and the system was solvated with the TIP3P water model[71] and neutralized with sodium ions. The resulting system contained approximately 870 lipid molecules in the bilayer and 470000 atoms in total. The system was first equilibrated in several 125-500ps steps during which harmonic restraints on the protein and membrane lipids were gradually removed.

After equilibrating the fast degrees of freedom, production simulations were done with the Hamiltonian Replica Exchange Molecular Dynamics (HREMD) method across 28 effective temperatures in a geometric progression from 300 to 420 K where exclusively protein atoms were treated with the Hamiltonian scaling factor [72–78] lambda, defined as 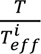 where *T* is the thermostat temperature and 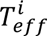 is the effective temperature of replica *i*. The very high temperature range promotes crossing of energy barriers to enhance conformational sampling and can also capture correlated contact responses that are not revealed at low temperatures. The Plumed HREMD implementation scales electrostatics (sqrt(lambda)), dihedral potentials, and LJ epsilon values. The protein, membrane, and solvent were separately coupled to a Nose-Hoover thermostat at a 1ps interval and a temperature of 300K. Pressure was controlled semi-isotropically with a Parrinello-Rahman barostat at a 5ps interval and a reference pressure of 1 bar. The Particle Mesh Ewald method was used for electrostatic calculations with a 1.2nm cutoff. The HREMD simulations were run using Gromacs 2019.6 patched with Plumed 2.6.1 [72,73,76,77,79,80] on the Xsede [81] Expanse cluster using a 2fs timestep integrated with the leap frog algorithm for 110 ns and replica exchanges attempted every 2ps. The exchange rate between neighboring systems averaged 15% +/-9% (standard deviation of 5% for all exchange pairs). All contact frequency based analyses were done using the averaged values, effectively quadrupling the enhanced sampling time per temperature to 440ns. Convergence was determined by observing little to no change in contact frequencies among most temperature-sensitive contacts at sensitization temperature (54 °C system) for the first two PCs (highest explained variance components) across the latter 50% of the simulation (Fig. S10).

### Contacts

After removing the solvent and lipids from each of the 28 systems we calculated all of the residues’ pairwise contact frequencies using GetContacts [82] which takes into account each residue’s unique chemical character and assigns different cutoff distances and interaction angles defining a contact accordingly. The contact frequencies for each pairwise interaction quantify the amount of simulation time the residue pair spends in contact. Approximately 11,000 different contacts occur on each protomer (including inter-protomer contacts) with nearly half of these being extremely rare or only occurring at extreme temperature. Principal component analysis of the averaged contact frequency data was performed with SciKit-Learn as 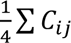 where *C_ij_* are identical contacts between *i* and *j* residues [83]. Analysis was done following the method described in Burns et al. [28]. The difference of roots test [29]was applied to determine the number of significant PCs for further analysis using 500 permutations of the original data to produce the null hypothesis contact set. The first 7 PCs were shown to be significant. The loading scores (eigenvector components) of the PCs were labeled with their initial variables (contact ids) and sorted according to their normalized absolute values to find the most temperature sensitive contacts on the given PCs. The top 18 (See SI Table 1) of the first 7 PC’s sorted loading scores were depicted on the structure using the contact’s PC on which it had its highest overall normalized loading score. The contact frequencies were projected onto each PC and plotted as a function of temperature and smoothed via spline interpolation.

### Network Analysis

The (inverse) contact frequencies were utilized as network edge weights with the NetworkX 2.8.4[84] library. Contacts that did not exceed 5% contact frequency in any of the natural temperature simulations (<= 60°C) were not included in the network calculations. The Girvan-Newman algorithm was used to deconstruct the averaged contact networks into 6 communities for the systems corresponding to 27°C and 54°C. The data from the averaged protomer was then applied to all four subunits to recreate the tetramer’s network from the averaged data for the edge betweenness centrality analysis. The same edge weights were used with the NetworkX edge betweenness centrality algorithm and the top 600 edges from the two temperatures were visualized on the structure as these accounted for essentially all the edges above the flatlined region of the sorted centralities (See Fig 4B).

### Cartesian Coordinate PCA

We separated the 54°C trajectory into the individual subunits, concatenated them into an extended single-subunit trajectory, and performed PCA to extract the motions relevant to activity at enhanced temperature. The MDAnalysis [85,86] implementation of PCA was used to calculate the principal components from the CA coordinates. For visualization, the all-atom trajectories were projected onto the PCs using the MDAnalysis method of applying a displacement vector to all the atoms in the residue corresponding to the CA atom used in the calculation of the PCs. From the frames where the structure exhibited a high RMSD value in the we extracted frames to represent the endpoints of the concerted motion. Residue RMSF was calculated from the CA coordinates of these 2 frames with MDAnalaysis and colors were applied along a gradient according to each CA’’s normalized RMSF (RMSF(resi)/RMSF(max)).

### Gate Measurements

The trajectories were aligned so that the pore was in the same position for each Hole calculation. The MDAnalysis implementation of Hole2 [56,87] was applied to each frame of each temperature’s trajectory and the minimum value from the coordinate range corresponding to the gate region was extracted.

### Random Forest

The frame-by-frame contact records from the systems ranging in temperature from 27°C to 58°C were concatenated. These temperatures were selected as they are all within temperatures where experimental activity has been recorded and at the peak of the *P*(*expanded gate*) and should therefore capture the contacts that are most physically relevant to the expanded gate. Each subunit’s set of contacts were separated and treated as individual samples. This effectively reduced the number of features per sample by ¾ to around approximately 6,300 features and quadrupled the total samples to approximately 300,000.

Contacts lining the pore were not included as they would directly inform on the state of the gate. To discretize the expanded and closed gate states and improve the accuracy of the model, we removed frames where the gate radius was between 0.75 and 1 Angstrom. The number of frames with a closed gate state greatly outnumbered the frames with the expanded state (90/10 ratio), so we applied the balanced subsample method to weight the two classes. We utilized results from two models trained with 3-fold cross validation and an 80/20 train/test split using 300 features and 700 estimators and 80 features and 100 estimators respectively. Both models produced nearly identical accuracies (88% precision and 64.5% recall for the 700 estimators and 88.5% precision and 64% recall for the 100 estimators). The contacts with feature importance above .001 (approximately 130 contacts) were retained as the feature importance values flatlined beyond this point (Fig 6A). The 106 contacts that were shared by both models’ top contacts were visualized on the structure and colored according to the PC on which they have their highest loading score.

### Visualizations

Visualizations were produced with PyMol and in-house scripts available on Github.

## ACKNOWLEDGMENTS

This work was supported by funds from the National Institute of General Medical Sciences with grant no. R35GM133488 (to V.V.) and grant no. R35 GM138243 (to D.P.).

## Notes

### Competing Interest Statement

The authors have declared no competing interest.

### Summary of Updates

Updated text and Figure 2

