## Supplemental Information for "Temperature Sensitive Contact Modes Allosterically Gate TRPV3"

TRPV3 Averaged Pairwise Contact Frequencies

| Temperature (°C) | A:LEU:135-A:LEU:138 | A:ASN:273-A:MET:320 | A:MET:159-A:PHE:157 | A:ARG:225-A:GLU:224 | A:ASN:242-A:ASP:288 | A:LYS:434-A:THR:431 |
|---|---|---|---|---|---|---|
| 27 | 0.942 | 0.506 | 0.999 | 0.849 | 0.544 | 0.860 |
| 30 | 0.942 | 0.508 | 1.000 | 0.863 | 0.590 | 0.861 |
| 34 | 0.939 | 0.503 | 0.999 | 0.880 | 0.587 | 0.860 |
| 38 | 0.939 | 0.502 | 1.000 | 0.883 | 0.585 | 0.859 |
| 42 | 0.936 | 0.505 | 1.000 | 0.868 | 0.589 | 0.868 |
| 46 | 0.933 | 0.499 | 0.999 | 0.858 | 0.593 | 0.865 |
| 50 | 0.935 | 0.498 | 1.000 | 0.838 | 0.603 | 0.870 |
| 54 | 0.936 | 0.503 | 0.999 | 0.840 | 0.610 | 0.871 |
| 58 | 0.935 | 0.506 | 1.000 | 0.845 | 0.622 | 0.865 |
| 62 | 0.933 | 0.510 | 0.999 | 0.845 | 0.618 | 0.862 |
| 66 | 0.935 | 0.508 | 1.000 | 0.841 | 0.624 | 0.858 |
| 71 | 0.935 | 0.489 | 0.999 | 0.809 | 0.580 | 0.858 |
| 75 | 0.936 | 0.484 | 0.999 | 0.788 | 0.565 | 0.861 |
| 79 | 0.934 | 0.493 | 0.999 | 0.794 | 0.554 | 0.864 |

Contact Frequency Principal Components

| | PC1 | PC2 | PC3 | PC4 | PC5 |
|---|---|---|---|---|---|
| A:GLU:263-A:SER:294 | 1.000 | 0.081 | 0.395 | 0.181 | 0.142 |
| A:ASP:293-A:TYR:260 | 0.982 | 0.032 | 0.725 | 0.241 | 0.314 |
| A:ASP:293-A:GLU:263 | 0.980 | 0.027 | 0.588 | 0.188 | 0.274 |
| A:GLN:292-A:HSD:244 | 0.934 | 0.381 | 0.375 | 0.132 | 0.038 |
| A:ARG:295-A:ASN:339 | 0.826 | 0.071 | 0.440 | 0.127 | 0.359 |
| A:ASN:297-A:ASN:338 | 0.821 | 0.105 | 0.376 | 0.035 | 0.264 |
| A:GLY:329-A:LEU:325 | 0.812 | 0.149 | 0.124 | 0.210 | 0.109 |
| A:LYS:246-A:SER:294 | 0.792 | 0.530 | 0.230 | 0.046 | 0.230 |
| A:ASN:297-A:ASP:293 | 0.783 | 0.122 | 0.469 | 0.297 | 0.312 |
| A:GLN:292-A:GLY:296 | 0.783 | 0.034 | 0.011 | 0.079 | 0.170 |
| A:LEU:281-A:THR:287 | 0.781 | 0.248 | 0.604 | 0.314 | 0.171 |
| A:GLN:313-A:VAL:306 | 0.757 | 0.292 | 0.146 | 0.098 | 0.401 |
| A:ASP:293-A:GLY:296 | 0.739 | 0.329 | 0.219 | 0.199 | 0.211 |
| A:ASN:242-A:GLN:286 | 0.737 | 0.085 | 0.435 | 0.094 | 0.335 |

S1. Top. Example contact frequency data. Bottom: PCA applied to the contact frequency data and sorted by PC1 loading score absolute values.

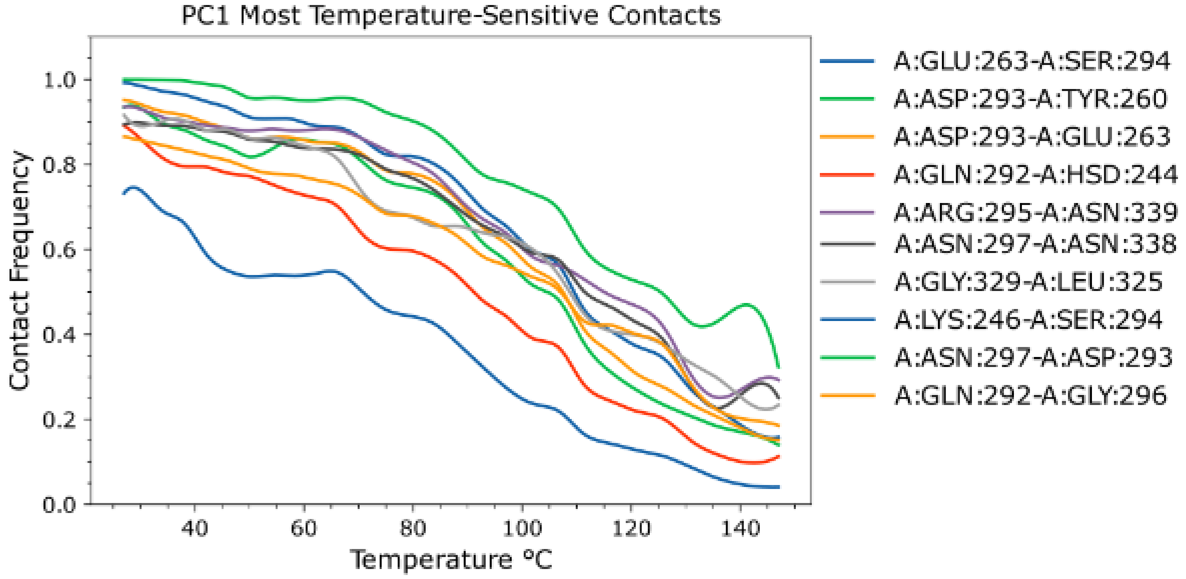
PC1 Most Temperature-Sensitive Contacts
Contact Frequency
Temperature °C
0.0
0.2
0.4
0.6
0.8
1.0
40
60
80
100
120
140
A:GLU:263-A:SER:294
A:ASP:293-A:TYR:260
A:ASP:293-A:GLU:263
A:GLN:292-A:HSD:244
A:ARG:295-A:ASN:339
A:ASN:297-A:ASN:338
A:GLY:329-A:LEU:325
A:LYS:246-A:SER:294
A:ASN:297-A:ASP:293
A:GLN:292-A:GLY:296

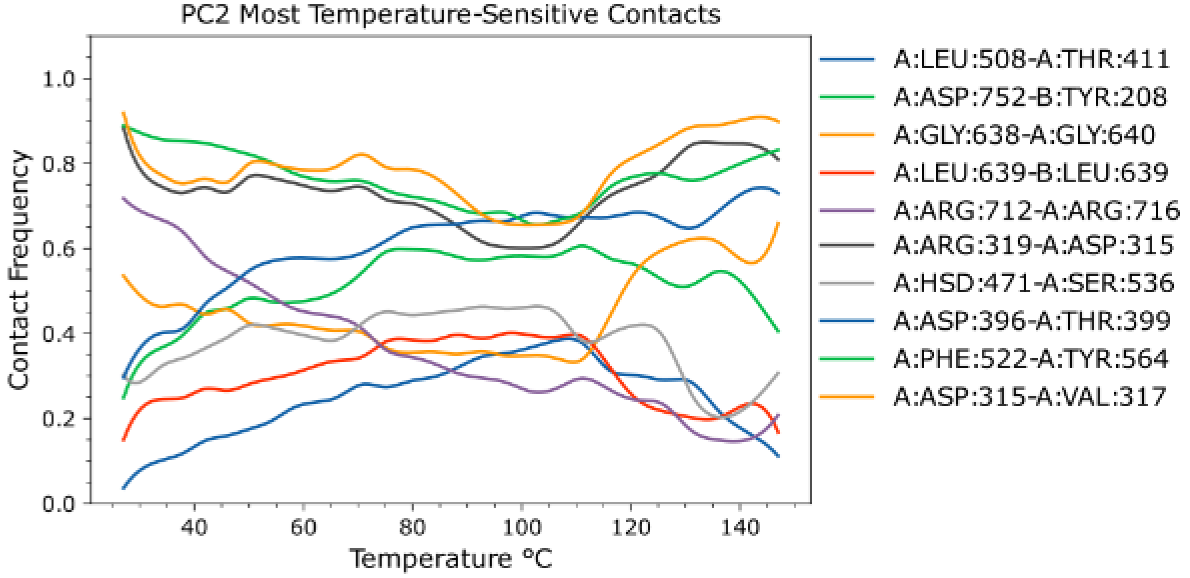
PC2 Most Temperature-Sensitive Contacts
Contact Frequency
Temperature °C
0.0
0.2
0.4
0.6
0.8
1.0
40
60
80
100
120
140
A:LEU:508-A:THR:411
A:ASP:752-B:TYR:208
A:GLY:638-A:GLY:640
A:LEU:639-B:LEU:639
A:ARG:712-A:ARG:716
A:ARG:319-A:ASP:315
A:HSD:471-A:SER:536
A:ASP:396-A:THR:399
A:PHE:522-A:TYR:564
A:ASP:315-A:VAL:317

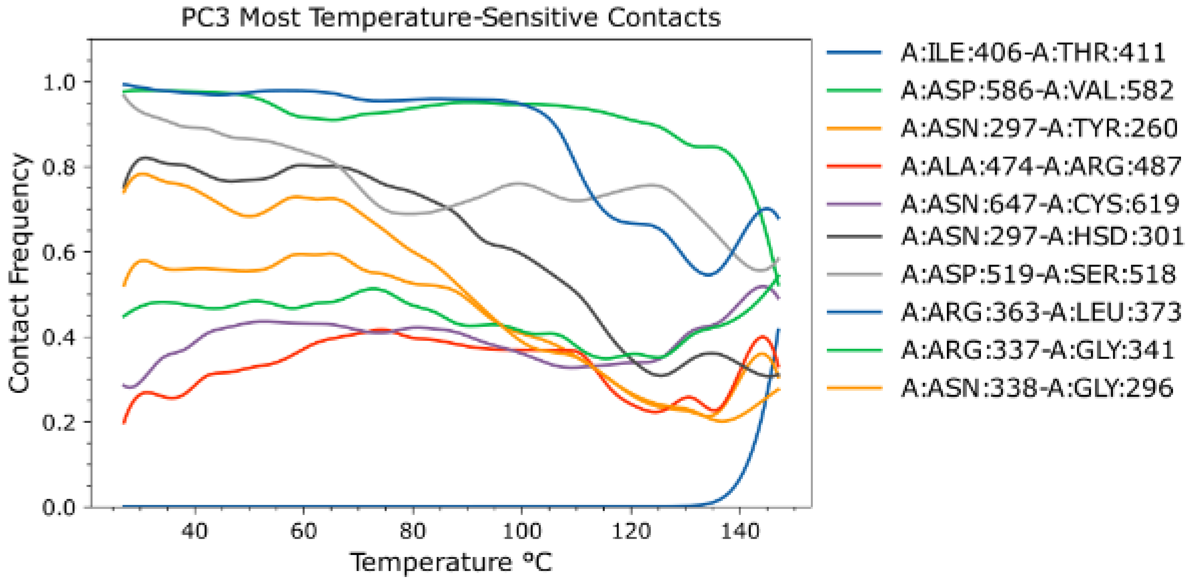
PC3 Most Temperature-Sensitive Contacts
Contact Frequency
Temperature °C
0.0
0.2
0.4
0.6
0.8
1.0
40
60
80
100
120
140
A:ILE:406-A:THR:411
A:ASP:586-A:VAL:582
A:ASN:297-A:TYR:260
A:ALA:474-A:ARG:487
A:ASN:647-A:CYS:619
A:ASN:297-A:HSD:301
A:ASP:519-A:SER:518
A:ARG:363-A:LEU:373
A:ARG:337-A:GLY:341
A:ASN:338-A:GLY:296

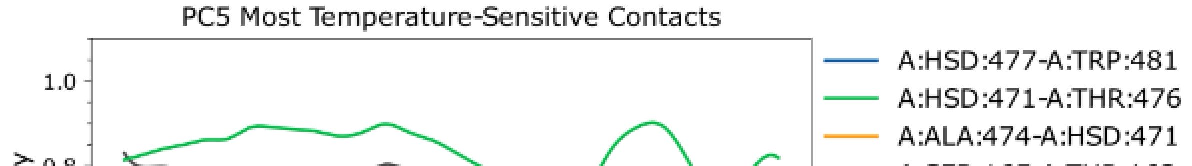
PC5 Most Temperature-Sensitive Contacts
1.0
A:HSD:477-A:TRP:481
A:HSD:471-A:THR:476
A:ALA:474-A:HSD:471

S2. Plots Top 10 most temperature-sensitive contact frequencies on the first 7 PCs. These correlate most closely with the corresponding PC projected trends.

| Rank | PC1 | PC2 | PC3 | PC4 | PC5 | PC6 | PC7 |
|---|---|---|---|---|---|---|---|
| 1 | A:GLU:263-A:SER:294 | A:LEU:508-A:THR:411 | A:ILE:406-A:THR:411 | A:LEU:730-A:THR:393 | A:HSD:477-A:TRP:481 | A:ALA:474-A:THR:476 | A:GLU:257-A:LYS:253 |
| 2 | A:ASP:293-A:TYR:260 | A:ASP:752-B:TYR:208 | A:ASP:586-A:VAL:582 | A:GLU:546-A:TYR:544 | A:HSD:471-A:THR:476 | A:ASP:315-A:VAL:317 | A:PRO:252-A:TYR:254 |
| 3 | A:ASP:293-A:GLU:263 | A:GLY:638-A:GLY:640 | A:ASN:297-A:TYR:260 | A:GLN:646-A:SER:621 | A:ALA:474-A:HSD:471 | A:ARG:567-A:THR:699 | A:VAL:737-B:HSD:256 |
| 4 | A:GLN:292-A:HSD:244 | A:LEU:639-B:LEU:639 | A:ALA:474-A:ARG:487 | A:ALA:164-A:ASP:166 | A:SER:165-A:THR:163 | A:ASP:586-A:THR:680 | A:ARG:729-A:SER:374 |
| 5 | A:ARG:295-A:ASN:339 | A:ARG:712-A:ARG:716 | A:ASN:647-A:CYS:619 | A:ASP:466-A:ASP:468 | A:HSD:301-A:TYR:260 | A:ALA:164-A:ASP:166 | A:ARG:371-A:ARG:729 |
| 6 | A:ASN:297-A:ASN:338 | A:ARG:319-A:ASP:315 | A:ASN:297-A:HSD:301 | A:GLY:168-A:SER:165 | A:GLU:465-A:GLU:467 | A:HSD:477-A:PRO:470 | A:ARG:226-A:ASN:273 |
| 7 | A:GLY:329-A:LEU:325 | A:HSD:471-A:SER:536 | A:ASP:519-A:SER:518 | A:ARG:729-A:SER:374 | A:ALA:474-A:TYR:540 | A:GLN:695-A:LEU:517 | A:ASN:251-A:GLY:247 |
| 8 | A:LYS:246-A:SER:294 | A:ASP:396-A:THR:399 | A:ARG:363-A:LEU:373 | A:ARG:729-A:THR:393 | A:ARG:487-A:LEU:491 | A:HSD:471-A:TYR:547 | A:TRP:742-B:ASN:273 |
| 9 | A:ASN:297-A:ASP:293 | A:PHE:522-A:TYR:564 | A:ARG:337-A:GLY:341 | A:ASN:330-A:SER:328 | A:HSD:471-A:TYR:547 | A:ASN:242-A:GLN:286 | A:LYS:353-A:VAL:304 |
| 10 | A:GLN:292-A:GLY:296 | A:ASP:315-A:VAL:317 | A:ASN:338-A:GLY:296 | A:GLN:646-A:ILE:609 | A:ARG:464-A:ASP:466 | A:LEU:473-A:TYR:540 | A:ASN:394-A:VAL:403 |
| 11 | A:LEU:281-A:THR:287 | A:GLY:638-B:GLY:638 | A:GLU:418-A:LYS:358 | A:LEU:473-A:LEU:475 | A:ALA:474-A:PRO:472 | A:GLU:682-A:GLY:678 | A:ARG:487-A:TRP:481 |
| 12 | A:GLN:313-A:VAL:306 | A:ARG:416-A:ASP:519 | A:GLU:334-A:TYR:321 | A:ARG:375-A:GLU:334 | A:ALA:543-A:LYS:545 | A:GLU:689-A:SER:685 | A:HSD:745-A:LYS:743 |
| 13 | A:ASP:293-A:GLY:296 | A:HSD:477-A:LEU:475 | A:ASP:586-A:LEU:584 | A:ASP:315-A:GLN:313 | A:LEU:482-A:SER:480 | A:GLN:274-A:ILE:223 | A:ASN:251-A:LYS:246 |
| 14 | A:ASN:242-A:GLN:286 | A:ALA:307-A:LYS:353 | A:TYR:382-B:GLN:216 | A:LEU:484-A:TRP:481 | A:ASN:643-A:GLN:646 | A:CYS:550-A:GLU:546 | A:HSD:477-A:LEU:475 |
| 15 | A:ASN:338-A:GLY:296 | A:ASN:338-A:HSD:301 | A:GLN:646-A:SER:621 | A:GLN:646-A:TYR:622 | A:ARG:150-A:LEU:152 | A:ARG:729-A:SER:372 | A:ALA:218-A:THR:217 |
| 16 | A:HSD:160-A:LYS:120 | A:GLU:418-A:LYS:358 | A:ASP:519-A:HSD:523 | A:PRO:755-B:PHE:249 | A:ASN:394-A:VAL:403 | A:ASN:683-A:GLU:687 | A:CYS:550-A:GLU:546 |
| 17 | A:ASN:297-A:GLN:292 | A:ASP:153-A:GLY:151 | A:ARG:149-A:VAL:154 | A:GLY:754-B:TYR:213 | A:GLY:486-A:VAL:490 | A:ARG:567-A:ARG:696 | A:ASN:735-B:HSD:256 |
| 18 | A:LYS:120-A:THR:163 | A:ASN:401-A:GLU:405 | A:ALA:206-A:ALA:218 | A:GLN:313-A:PHE:310 | A:GLU:308-A:PHE:310 | A:MET:479-A:TRP:481 | A:ASN:394-A:SER:402 |

S3. Top 99.85th percentile of temperature-sensitive contacts for the first 7 contact PCs.

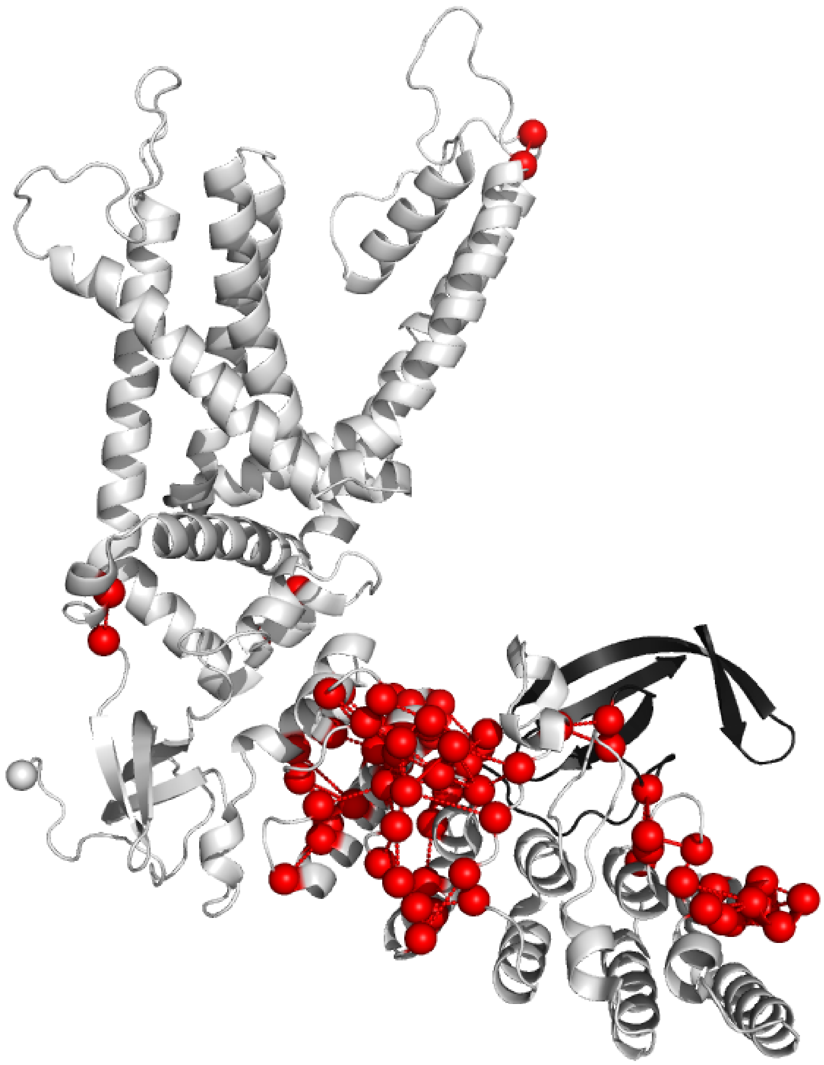

S4. 99.5th percentile of PC1 contacts showing the heavy clustering around ankyrin repeat 5. Black beta strands are the IPCD of the adjacent protomer

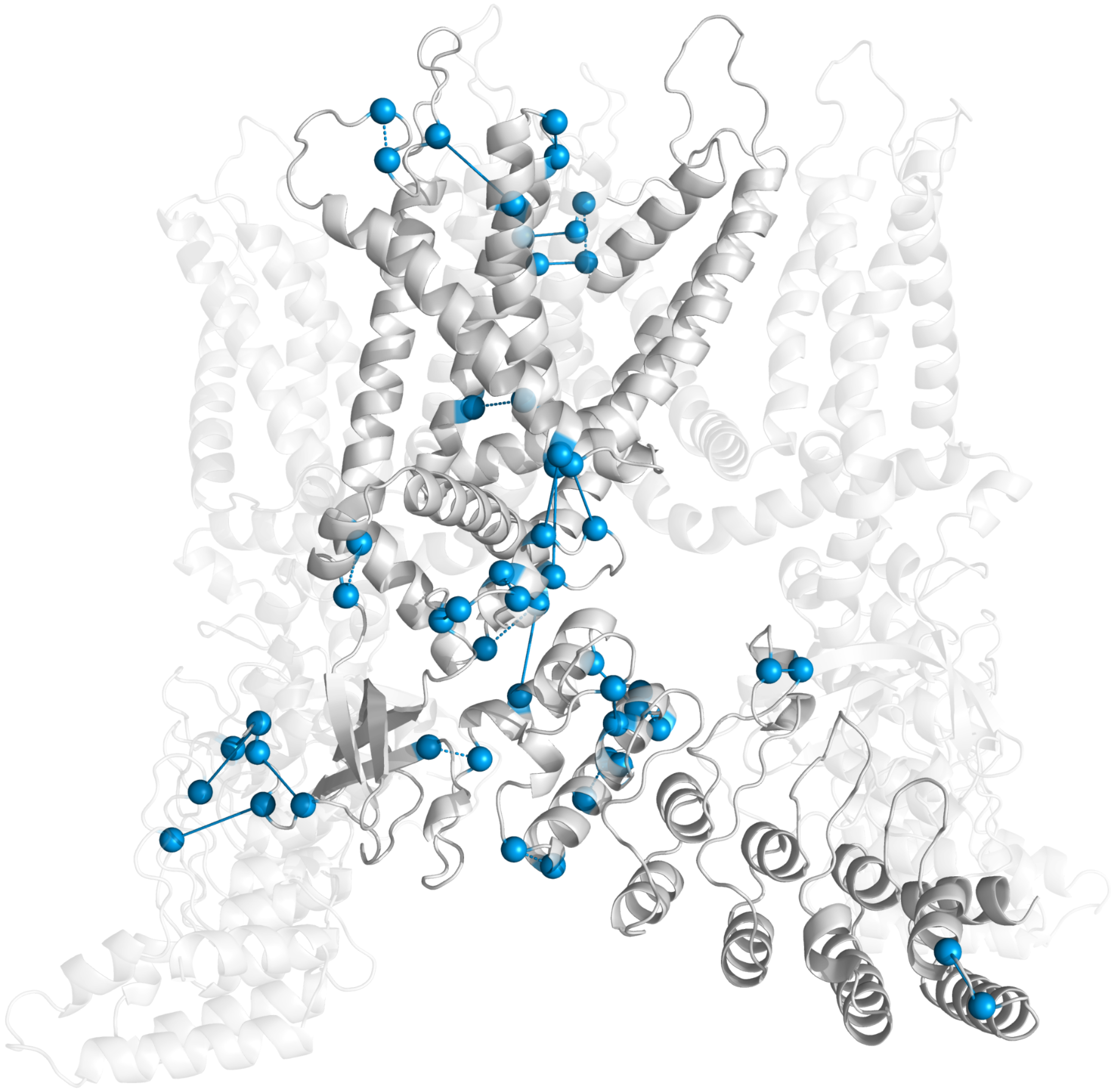

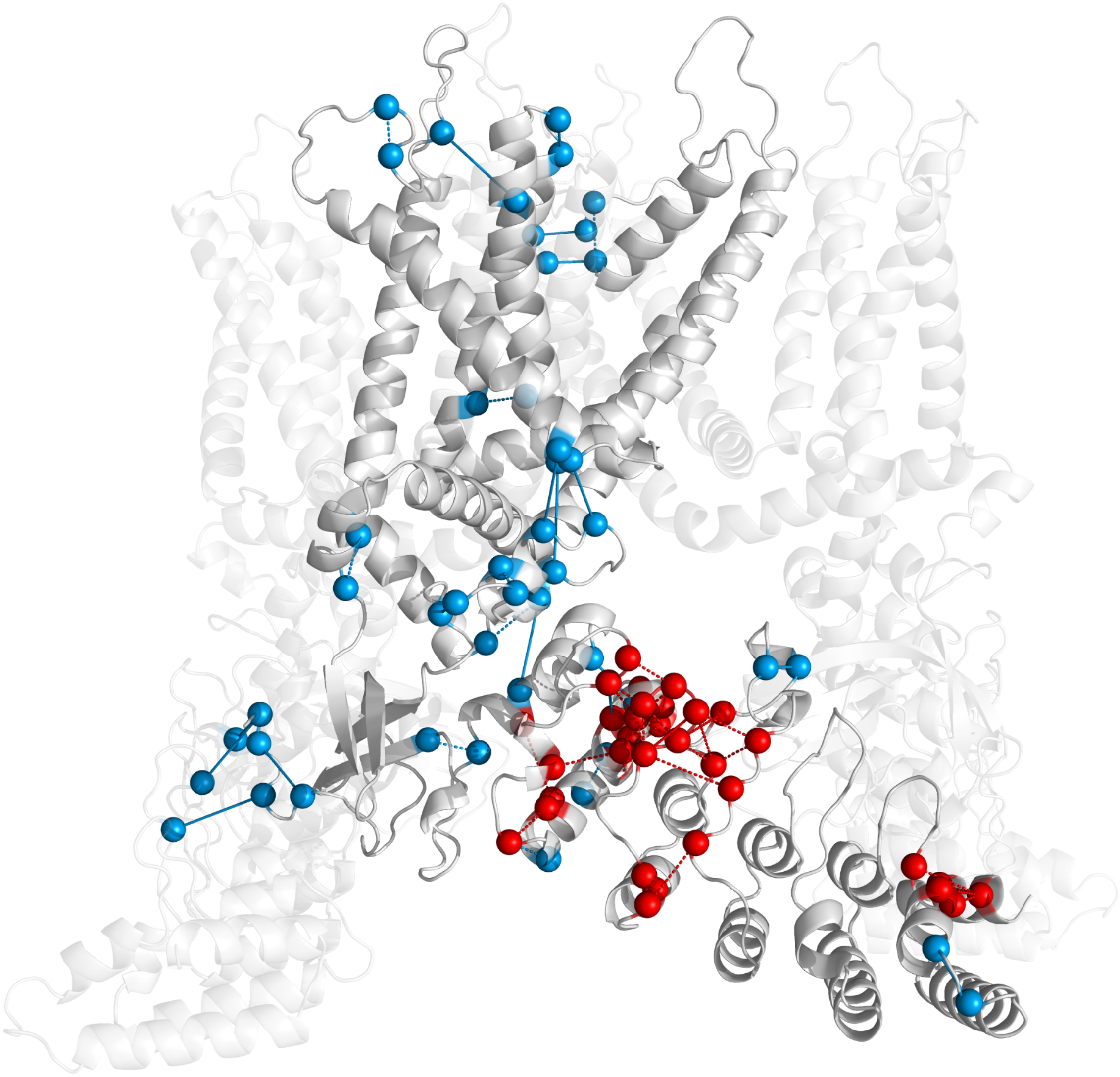

S5. Top: 99.75th percentile of PC2 contacts. Bottom: Top 99.75th percentile of PC1 and PC2. Disorder (red/ dashed lines) develops on the upper ARD while order (blue/ solid lines) increases around the center of the protomer with temperature.

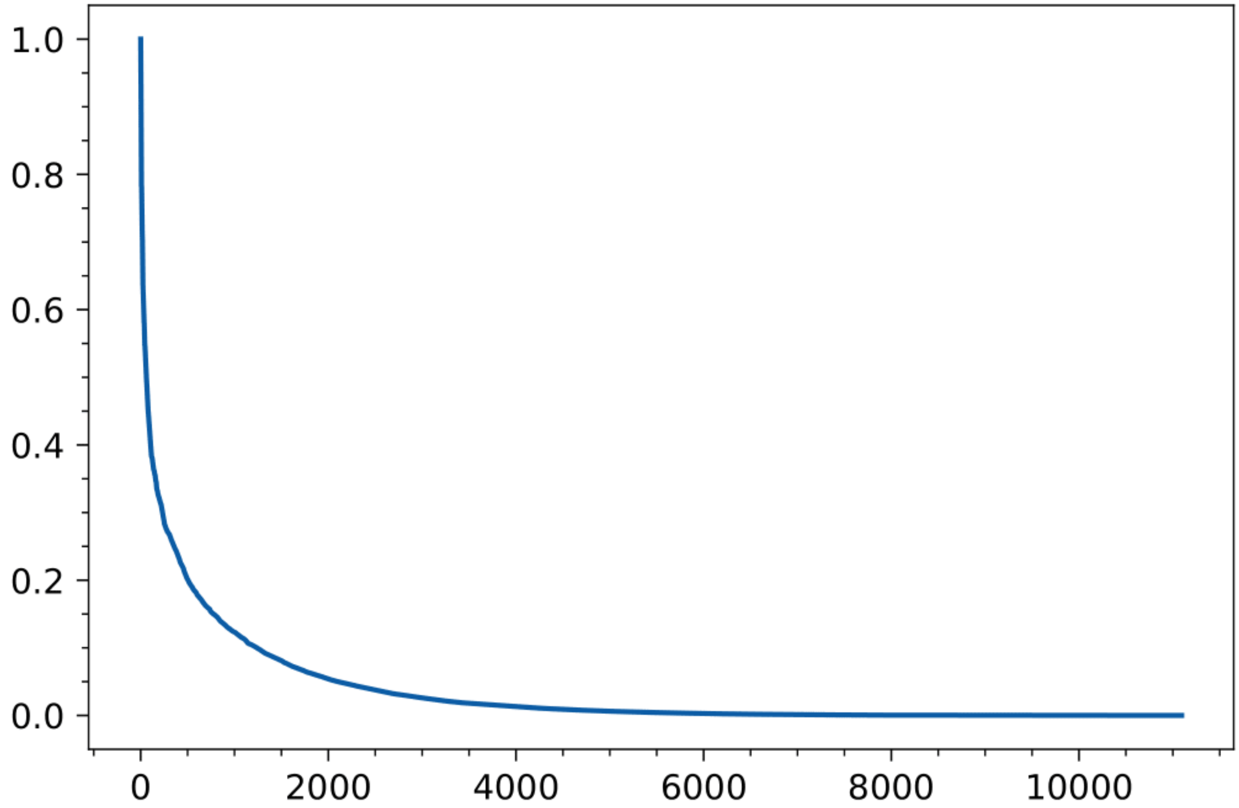


S6. Loading scores decay. Each PC exhibits a similar rapid drop in loading scores.The top 99.85th percentile is populated by contacts with loading scores no lower than 0.55 on any of the first 7 PCs.

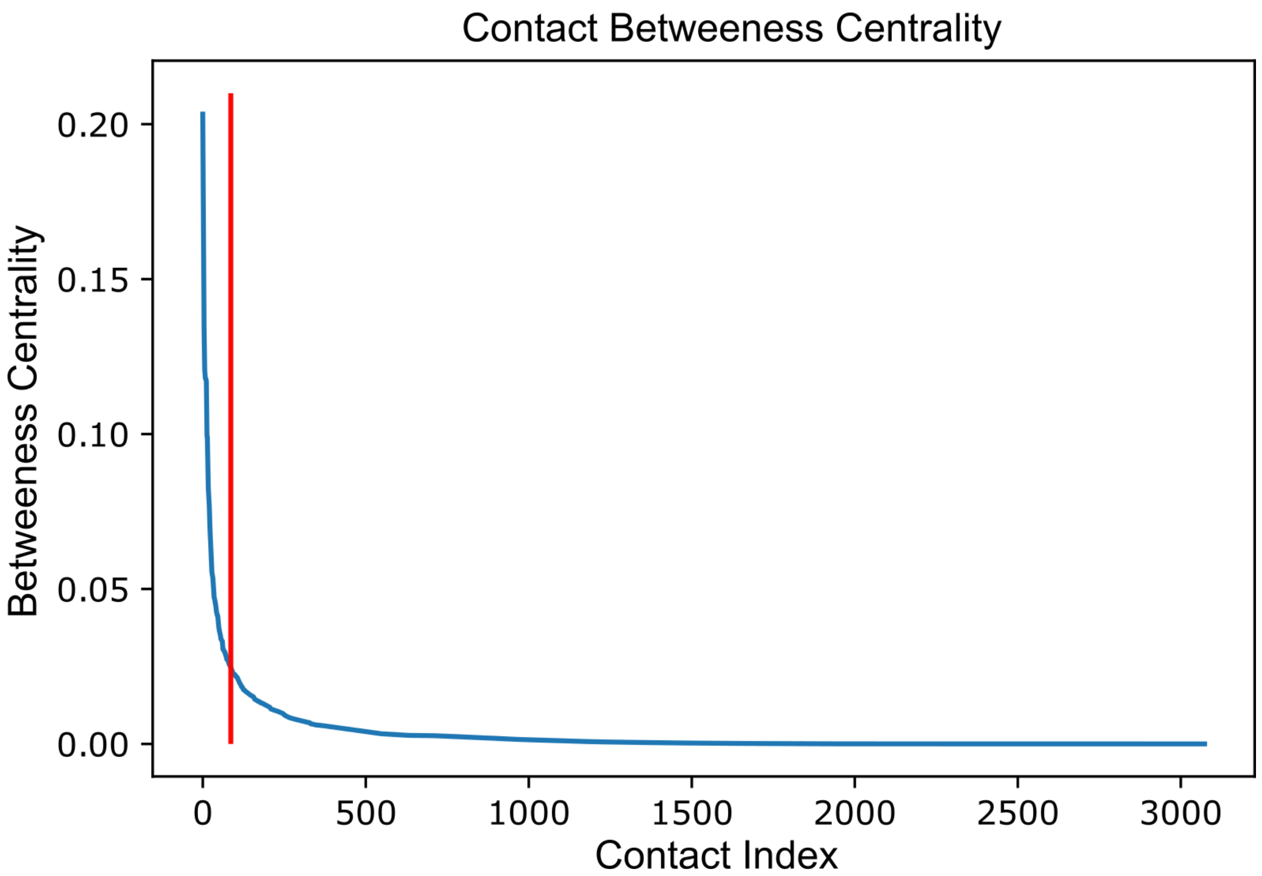

S7. Betweenness-centrality in descending order with the highest betweenness edges that constitute the “allosteric skeleton” occurring left of the vertical red cutoff bar.

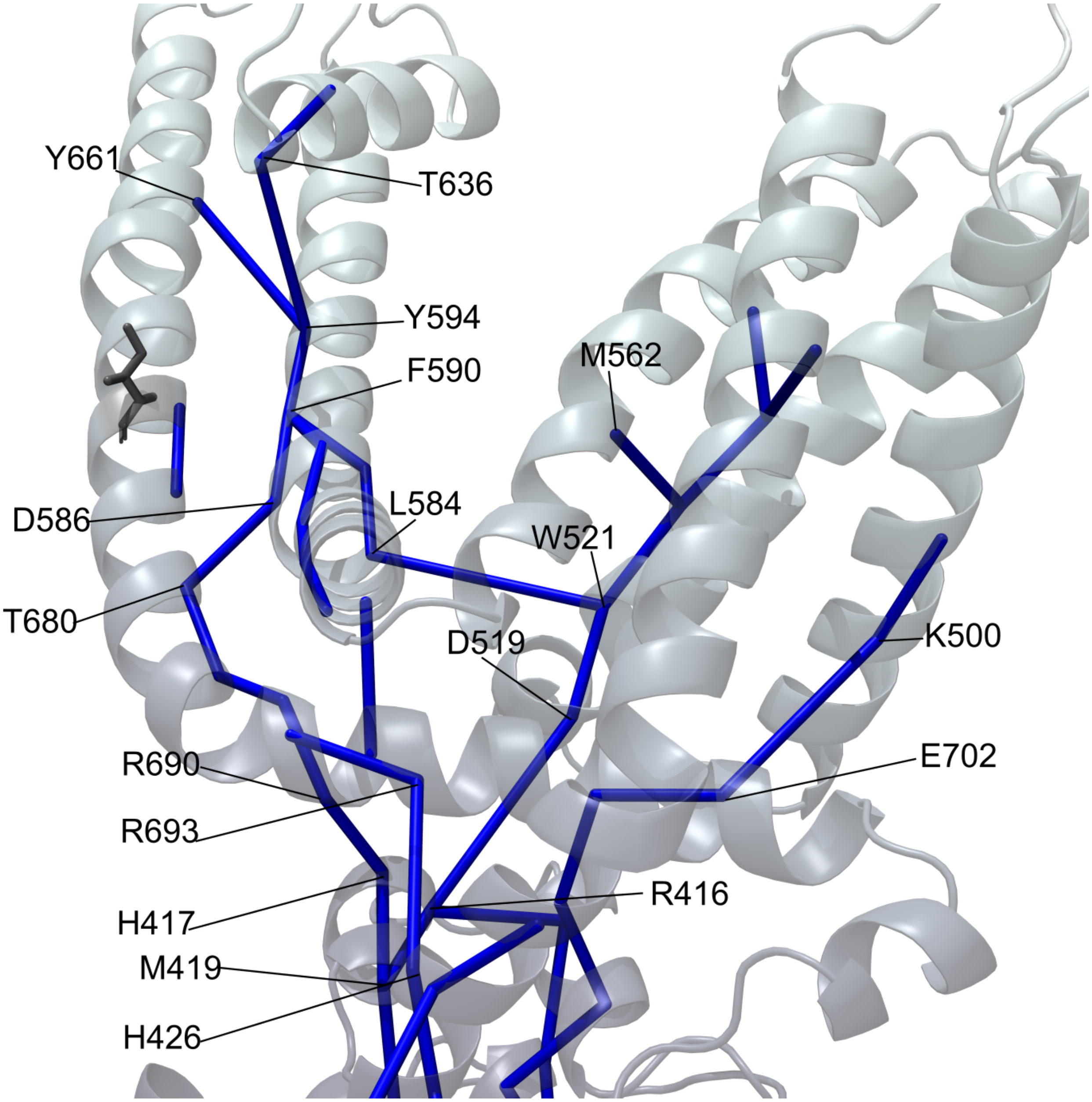


S8. Close up of the transmembrane allosteric skeleton. (Need to finish labeling this)

S9.

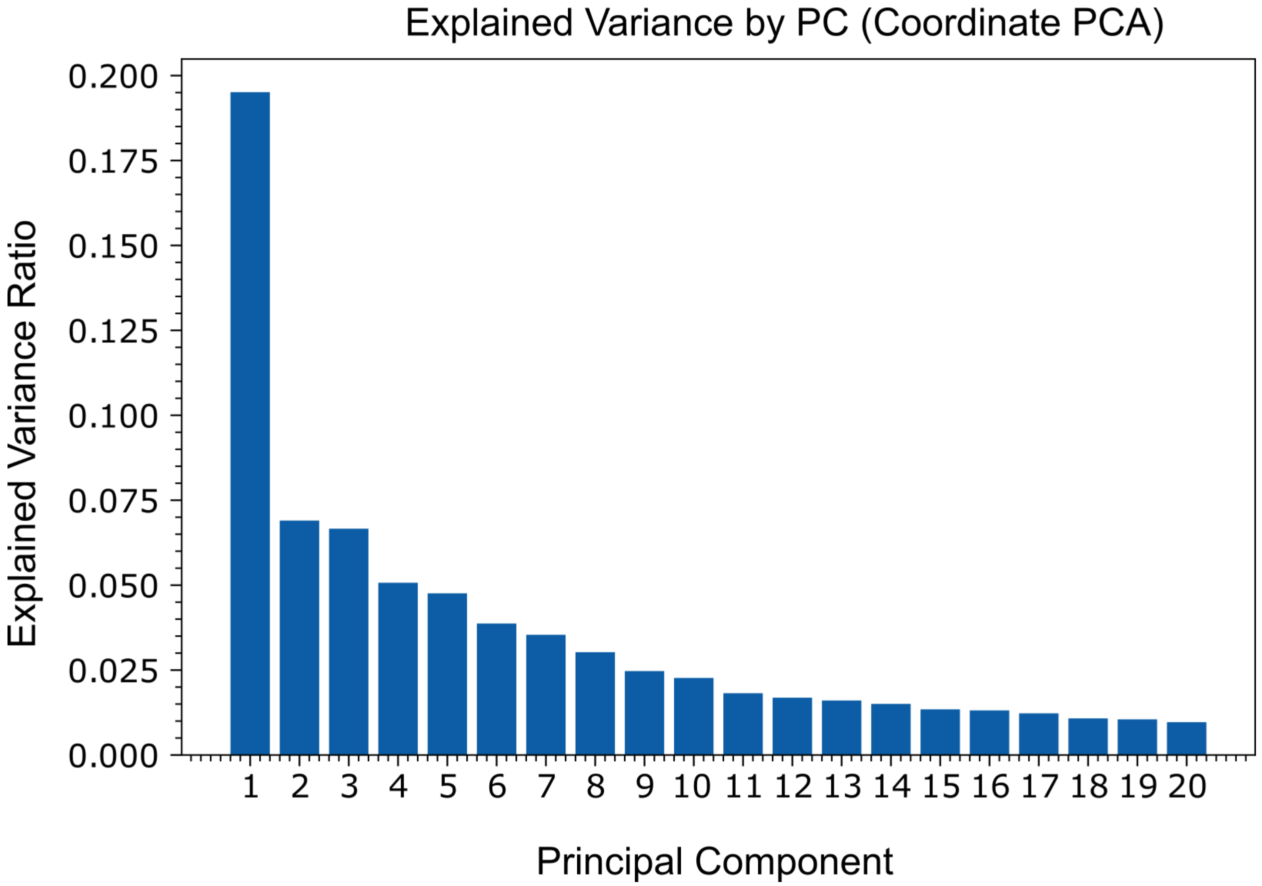


S9. Explained variance of the cartesian PCs.

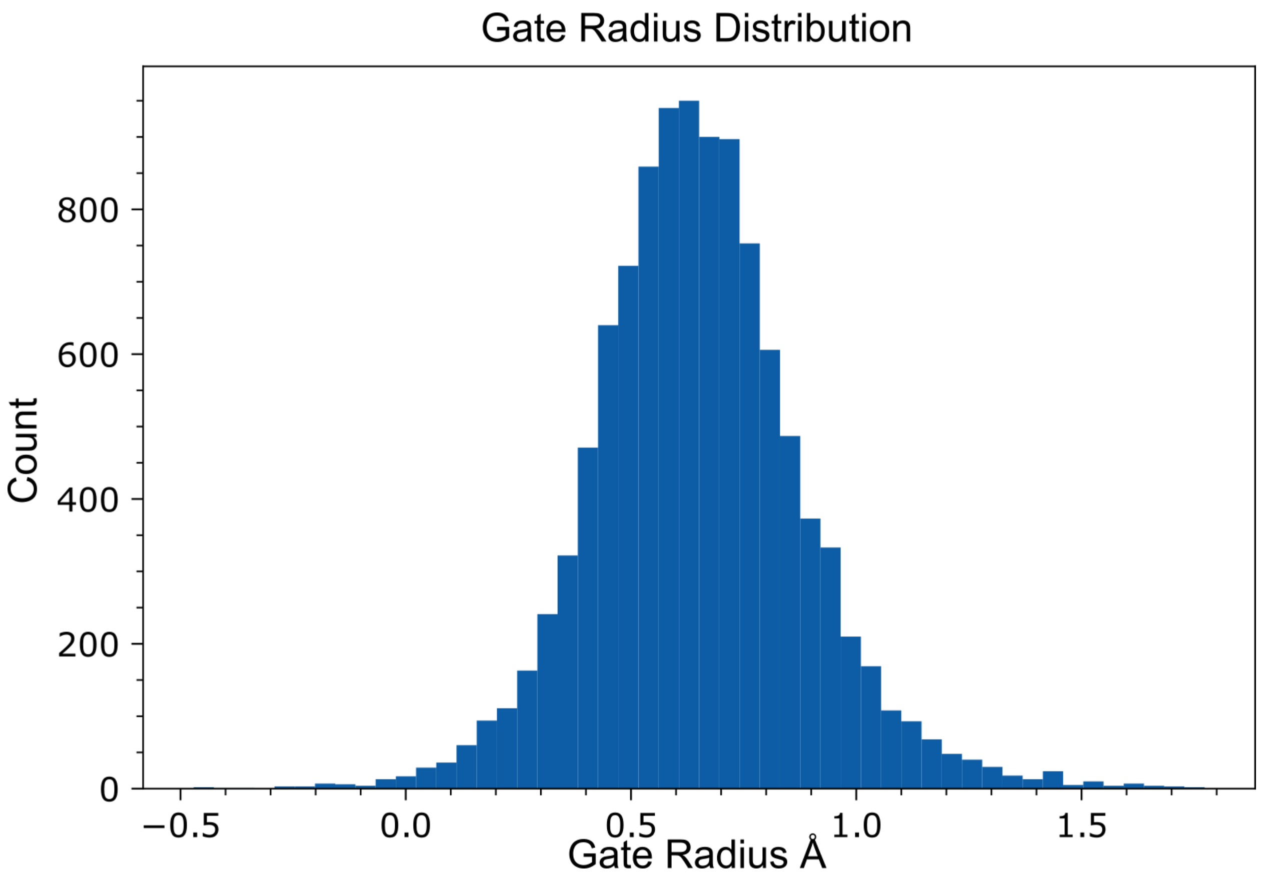

S10. Distribution of the gate radius within the physiological temperature range.

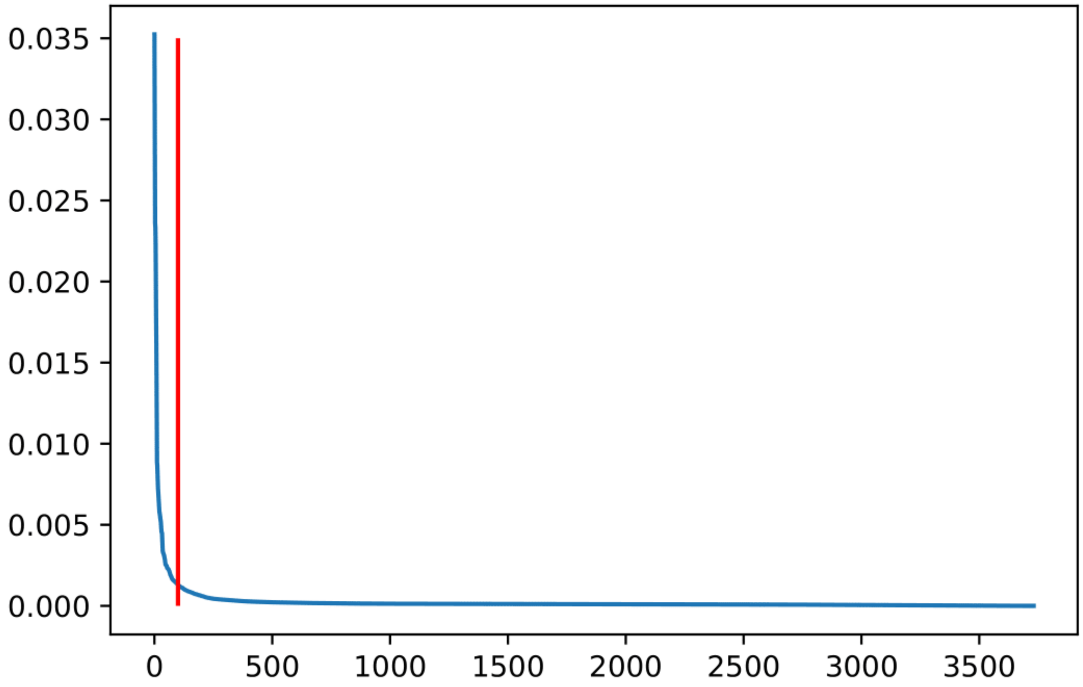


S11. Decay in feature importance from a random forest model. The top 100 feature important contacts were chosen (left of vertical red cutoff bar) because the remaining contacts have values of essentially 0.

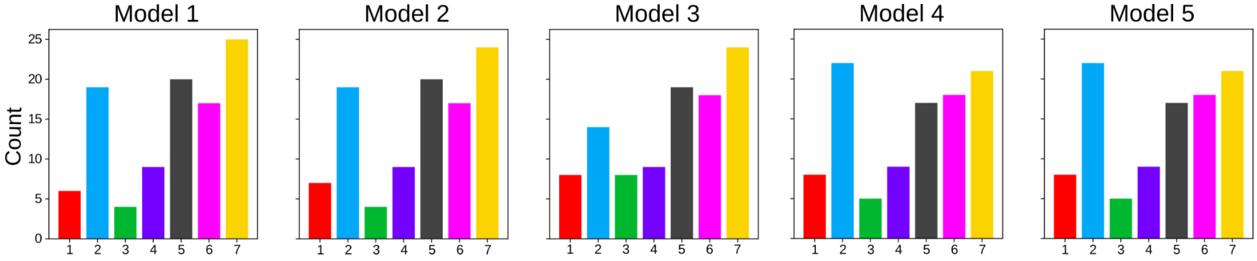


S11. Distribution of the top 100 feature important contacts among the contact PCs that they score highest on for the 5 random forest models. Models 2 and 5 are considered general models and were used for further analysis.

Model 2

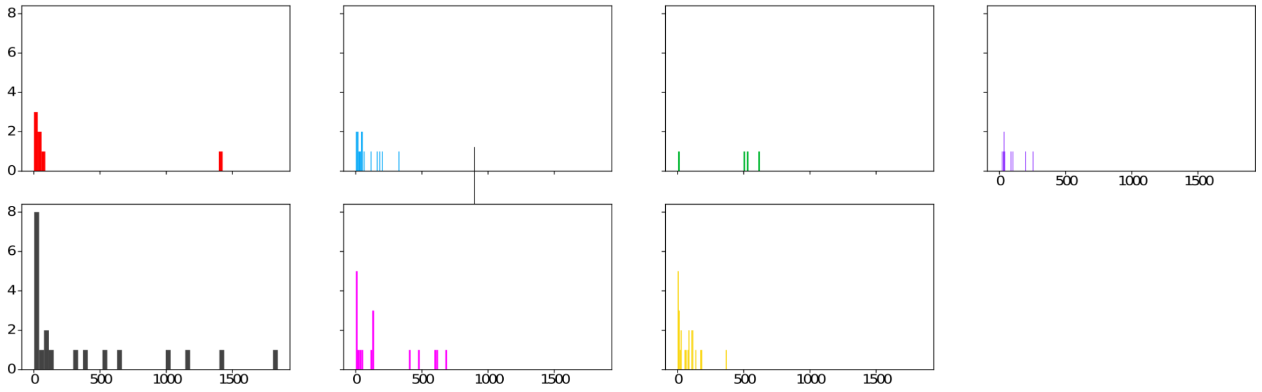


Model 5

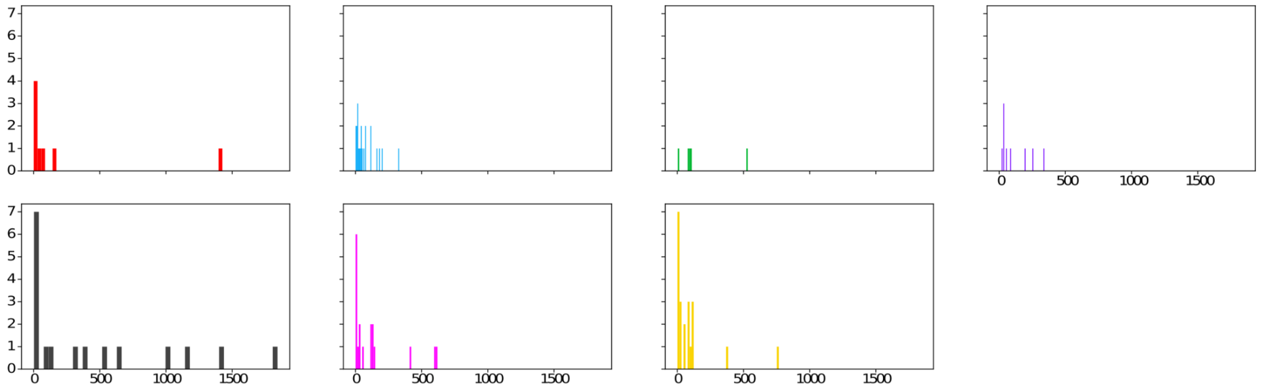


S12. The Temperature sensitivity rank histograms by PC taken from the top 100 feature important contacts from models 2 and 5. Left on the x-axis corresponds to highly temperature-sensitive contacts.

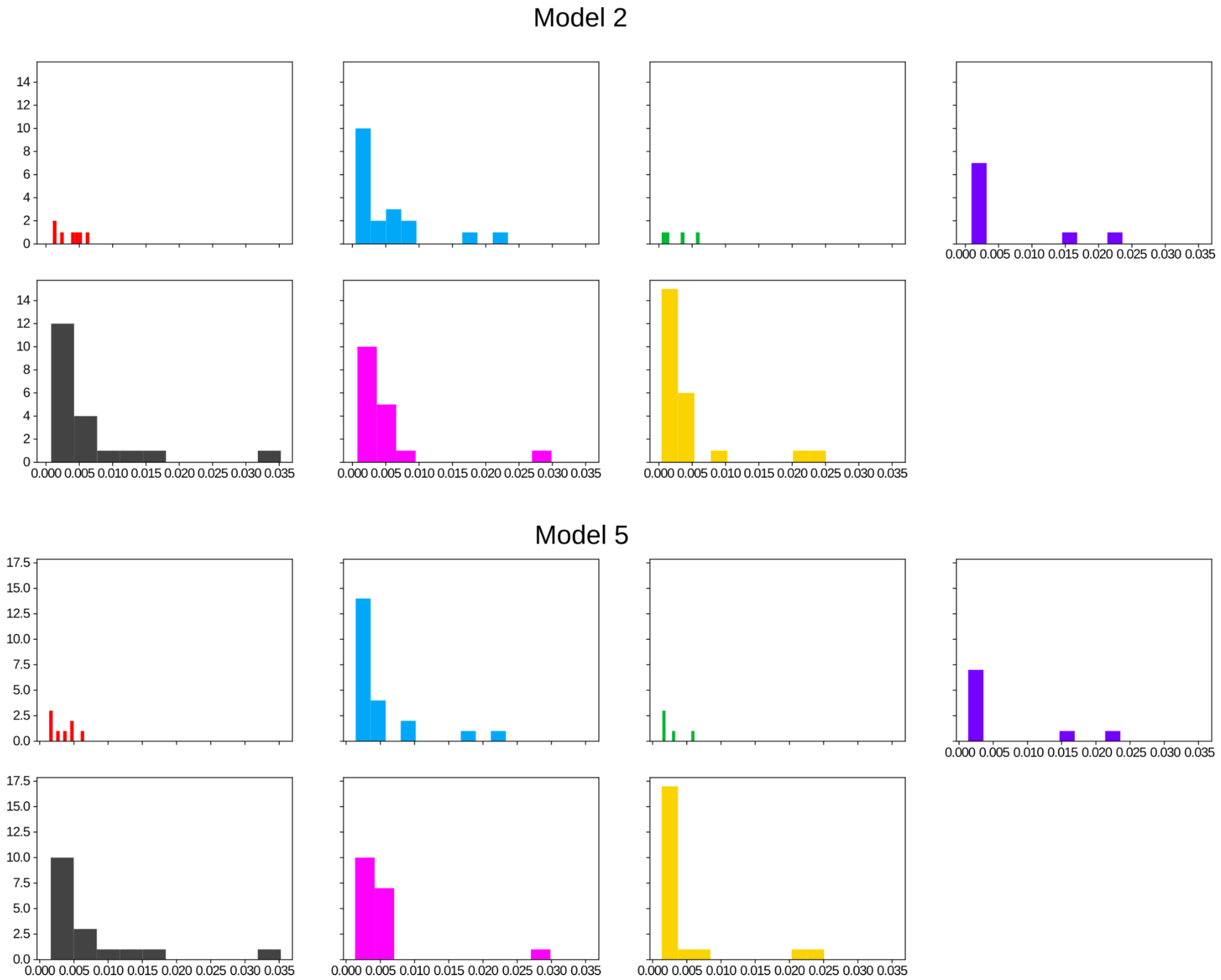


S13. Histograms of feature importance values for the top 100 contacts from models 2 and 5 according to the pc that the contacts have their highest loading scores on. Right on the x-axis corresponds to high feature importance.

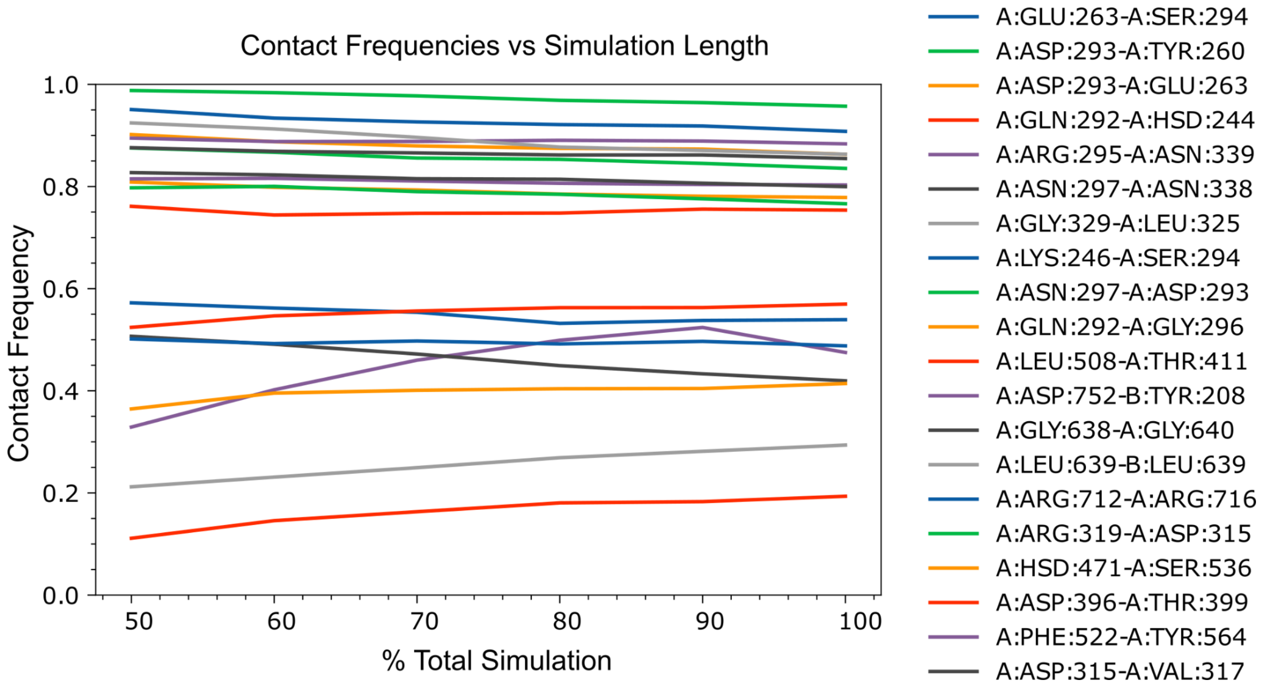


S14. Convergence of PC1 and PC2’s most temperature-sensitive contact frequencies from the 54 C simulation.
